# A time-calibrated phylogeny of hummingbirds supports stepwise diversification in the Andes

**DOI:** 10.64898/2026.02.06.704360

**Authors:** Rute R. da Fonseca, Gary R. Graves, Siavash Mirarab, Inger Winkelmann, Santiago Claramunt, Miguel M. Fonseca, Jose A. Samaniego Castruita, Fernando Penaloza, Cai Li, Sara Rocha, Merly Escalona, Lys Sanz Moreta, Anders Albrechtsen, Tandy Warnow, Jon Fjeldså, M. Thomas P. Gilbert, Carsten Rahbek

## Abstract

Hummingbirds (Trochilidae) represent the second largest avian family, with ∼356 species occupying diverse habitats across the Neotropical and Nearctic regions. Their extensive diversification includes notable adaptations to extreme environments, with nearly one-third of extant taxa belonging to the predominantly high-altitude Coquettes and Brilliants. Although recent work proposed a monophyletic Andean clade uniting these groups, consistent with rapid radiation during Andean orogeny, we find no support for this relationship. Using 2,949 nuclear loci sampled from 47 species spanning all nine major hummingbird clades, we recover a different evolutionary pattern: a stepwise sequence of diversification in which Brilliants are sister to a broader assemblage comprising Coquettes, the genus *Patagona*, Emeralds, Mountain Gems, and the recently diverged Bees. By assembling complete mitochondrial genomes, we additionally detect significant discordance between nuclear topologies and those derived from the mitogenome and Z chromosome. Analyses of gene-tree heterogeneity show that incomplete lineage sorting is pervasive across the phylogeny, with particularly strong impacts on branches associated with Andean diversification. Divergence-time estimation further indicates that the major Andean radiation—including Brilliants, Coquettes, Emeralds, Bees, and Mountain Gems—originated around ∼14 Ma, with the three younger clades diversifying ∼12 Ma, coinciding with both the mid-Miocene Andean uplift and the mid-Miocene Climate Transition that increased habitat heterogeneity and likely promoted rapid speciation. To support future phylogenomic efforts, we identify a reduced set of highly informative, independent protein-coding loci and present a near-complete species-level phylogeny constrained by our autosomal backbone. Our findings highlight the importance of integrating loci with distinct inheritance modes to detect and interpret phylogenetic incongruence in rapid radiations.

## Introduction

Hummingbirds (family Trochilidae) constitute one of the largest avian families (Howard 2003), resulting from a rapid radiation that began in the lowlands of South America in the Early Miocene (Bleiweiss 1998a). The tropical Andean region, which underwent a major uplift about 10 million years ago, provided new ecological niches that facilitated speciation. About 78% of hummingbird species belong to a monophyletic clade that may have originated in the Andes, and nearly 40% of the 356 extant species (Chesser et al., 2022; Remsen et al., 2023), currently inhabit the Andes mountains (McGuire et al. 2014; Rahbek and Graves 2000). Hummingbirds include the smallest birds in the world (several species with body masses of 2.0-2.3 g) and possess extreme physiological and morphological adaptations that facilitate demanding acrobatic displays and sustained hovering flight (Altshuler and Dudley 2002; Altshuler et al. 2004; Skandalis et al. 2017). Hummingbird species are traditionally divided into nine major clades, which are assumed to be monophyletic (Bleiweiss 1998b; McGuire et al. 2014): (i) Topazes (sister to the rest of Trochilidae), (ii) Hermits, (iii) Mangoes, (iv) Brilliants and (v) Coquettes (iv and v mostly Andean), (vi) Emeralds, (vii) the genus *Patagona* (the giant hummingbird complex that includes the largest hummingbird species), (viii) Mountain Gems, and (ix) Bees (Bleiweiss 1998b) (formal clade nomenclature from (McGuire et al. 2009)). We performed targeted sequencing of 2949 nuclear genes from 46 species (13.6% of recognized species) representing the nine major clades of hummingbirds. We have also reconstructed the complete mitochondrial genomes for all 46 species. We found that the nuclear and mitochondrial topologies (hereafter nuDNA and mtDNA, respectively) are discordant relative to the placement of Brilliants and Coquettes. Our nuDNA markers agree with Bleiweiss’s phylogeny (based on DNA-DNA hybridization distances (Bleiweiss 1998b)) that showed Brilliants and Coquettes as non-sister clades. In contrast, both mtDNA and sex chromosome Z phylogenies suggest that Brilliants and Coquettes are sister taxa, as in the McGuire et al. tree (built using 4 nuDNA and 2 mtDNA markers and dense sampling of 284 hummingbird species). The nuDNA data showed high gene tree discordance relative to the branch that separates Brilliants and the clade containing Coquettes, Emeralds, the genus *Patagona*, Mountain Gems, and Bees. This can be explained by incomplete lineage sorting, associated with a very rapid diversification associated with the short branch. In this scenario, our large data set and the separate analysis between loci with different effective population sizes and different inheritance modes were crucial in detecting the phylogenetic incongruence.

## Materials and methods

### Samples

Species were chosen so that all the major trochilid clades and subclades reported by Bleiweiss et al. (Bleiweiss et al. 1994) and McGuire et al. (McGuire et al. 2014) were represented. Tissue samples were obtained from the National Museum of Natural History, Smithsonian Institution (n = 33 species) and the Louisiana State University Museum of Natural Sciences (n = 13 species) (see Table S1 for details)

### Target selection and bait design

We selected one-to-one orthologs between chicken (*Gallus gallus*) and zebra finch (*Taeniopygia guttata*) as annotated in ENSEMBL version 66 annotation (more details in the Supplementary Methods) resulting in a final set of 166322 probes corresponding to 2949 genes and summing up to approximately 20 Mb (for an approximately 7 Mb of total captured sequence).

### DNA extraction and capture-enrichment

Details regarding the laboratory procedure can be found in the Supplementary Methods. To confirm that laboratory procedures worked correctly, we captured and sequenced the first eight species in Table S2 using 250bp paired-end sequencing on the Illumina MiSeq platform. Once this was confirmed, the remaining 38 samples were captured in batches of eight and sequenced on an Illumina HiSeq 2500, using one lane of 100bp paired-end sequencing per eight samples.

### *De novo* assembly of nuDNA capture regions and orthology determination

The adapters were removed with CUTADAPT (Martin 2011) and the reads were quality trimmed/filtered with PRINSEQ (Schmieder and Edwards 2011). The overlapping read pairs were merged with FLASH (Magoč and Salzberg 2011). SOAPdenovo (Luo et al. 2012) was used to *de novo* assemble the reads using a k-mer length of 63. The contigs for each sample were aligned against the reference exons and the best hit for each exon was selected using lastz (Harris 2007) and chainNet (https://genome.ucsc.edu/). Exons covering less than 50% of the reference were discarded. Coding regions were predicted with Exonerate (Slater and Birney 2005), and exons were translated and grouped per gene.

### Phylogenetic reconstruction of the nuDNA markers

Amino acid sequence alignments of the predicted coding regions were generated with MAFFT (Katoh 2002) (with the option L-INS-i for a more accurate alignment). Phylogenetic reconstructions were done using the corresponding back-translated nucleotide sequences. The nucleotide sequence for all the nuclear gene orthologs were concatenated for estimating a phylogenetic tree using the maximum likelihood program RAxML under the GTR+GAMMA model of sequence evolution. Trees were built using different partitions per codon position and three different gene subsets (Table S3). ASTRAL-II (Mirarab and Warnow 2015) was used for a genome-scale coalescent-based species tree estimation. Best ML gene trees were estimated using RAxML (Stamatakis 2014) version 8.0.20 and starting from 10 random starting trees, using the GTR+GAMMA mode. We also performed 200 replicates of bootstrapping per gene using RAxML (Stamatakis 2014). We used ASTRAL (Mirarab et al. 2014) version 4.10.0 to estimate the species trees from the best ML gene trees. We estimated support by using multi-locus bootstrapping using site-only or gene-site procedure and also ASTRAL’s built-in local posterior probabilities (Sayyari and Mirarab 2016). To assess gene tree discordance (Figure S1), we computed the number of gene trees that were incompatible with the species tree, first by using binary gene trees, and then after we contracted gene tree branches with support below 75%. We also quantified quartet support using ASTRAL’s built-in function that computes quartet frequencies around a branch according to gene trees (Figure S2). For each quartet defined around each branch, three possible topologies are possible; ASTRAL outputs the average frequency of the topology seen in the species tree and the two alternatives. Results were visualized using DiscoVista (Sayyari et al. 2017). A larger constrained phylogenetic tree including 284 species (see Data Availability) was constructed by extracting each clade’s subtree from a previously published phylogeny (McGuire et al. 2014) and grafting these subtrees into the 9-clade backbone obtained with our nuDNA markers.

### Assembly of full mtDNA genomes, annotation and sequence alignments

The MITObim (Gan et al. 2014) pipeline was used to reconstruct the full mtDNA genomes for the 46 hummingbirds using the mtDNA genome assembly of *Archilochus colubris* (NC_010094.1). Briefly, MIRA 4 (Chevreux 2005) was used for the initial mapping assembly followed by iterative mapping using MITObim.pl script for 10 iterations. In order to remove IUPAC ambiguity characters resulting from SNPs in the assembly, miraconvert was used to create a consensus from them. The reconstructed mtDNA genomes were initially annotated using MITOS (Bernt et al. 2013) and ARWEN (Laslett and Canbäck 2008). The gene sequences were aligned with MAFFT version 7 (Katoh and Standley 2013) with options G-INS-i (for codon and amino acid sequences) and L-INS-i (for rRNA sequences) with 1000 iterations, using the BLOSUM62 scoring matrix (Henikoff and Henikoff 1992) for the amino acid sequences (the codon sequences were also aligned using the translated amino acid sequences) and the default 200PAM/k=2 scoring matrix for the rRNAs sequences. The alignments were filtered using GUIDANCE version 1.5 (Penn et al. 2010) with the following parameters: 100 bootstrap replicates, column cutoff and site masking score of 0.8. The resulting single gene alignments were finally concatenated into 13,808 nucleotides (13 protein-coding genes plus 2 rRNA sequences) or 3,791 amino acids.

### Phylogenetic reconstruction of the full mtDNA genomes

The mtDNA genome tree was estimated using two concatenated alignment sets: (i) the concatenated nucleotide sequences for the 13 protein-coding genes and of the two ribosomal RNAs and (ii) the concatenated amino acid sequences of the 13 mtDNA protein-coding genes. The software RAxML (Stamatakis 2014) version 8.1.7 with 100 rapid bootstrap replicates and 20 ML searches was used to estimate the phylogenetic trees under the GTR+GAMMA and MtMam models of sequence evolution for the nucleotide and amino acid datasets, respectively. Trees were estimated using the best partitioning scheme proposed by the statistical approach “greedy” with branch lengths linked among partitions implemented in PartitionFinder or PartitionFinderProtein v1.1.1 (Lanfear et al. 2012). The initial partition blocks were defined per codon position for the nucleotide dataset (but not for the rRNA sequences) and per gene for the amino acid dataset and the rRNA genes.

### Phylogenetic discordance

Gene concordance factors (Minh et al 2020a) were computed in IQ-TREE (Minh et al 2020b) and used to quantify the proportion of loci supporting each internal branch of the reference phylogeny. Gene concordance factors provide a measure of gene-tree agreement across loci and explicitly characterize phylogenomic discordance arising from processes such as incomplete lineage sorting or introgression, independent of resampling-based node support. Furthermore, gene-tree clustering was examined using TreeCl, and the topological heterogeneity among loci was assessed by computing pairwise geodesic distances between gene trees in treespace (Kendall and Colijn 2016). These distances quantify the shortest path between trees under a metric that accounts for both topology and branch length differences. The resulting distance matrix was summarized using classical multidimensional scaling (MDS). Topological correspondence between mitochondrial and nuclear phylogenies was visualized using a tanglegram constructed in R (packages *ape, phytools, dendextend, phylogram, viridis*, and *dplyr*), enabling direct comparison of branching order and taxon placement between the two trees.

### Divergence time estimation

For molecular dating, we used a subset of 259 exons corresponding to nuclear gene sequences chosen according to the following criteria: i) GC content between 0.43 and 0.5 (first and third quartiles of GC distribution); iii) 90% of the length of the exon covered in a minimum of 40 species (maximum 10% missing sequence); iv) depth of coverage between 88 and 213 (first and third quartiles), calculated after mapping the probes to the reference genome of the Anna’s hummingbird (Zhang et al. 2014). Variants were called after joint genotyping using GATK version 4.0.7.0 (DePristo et al., 2011), and hard-filtered using the following options: “QD < 2.0 || FS > 60.0 || MQ < 40.0 || MQRankSum < -12.5 || ReadPosRankSum < -8.0 || SOR > 3.0”. Consensus sequences with ambiguity codes were extracted with bcftools (Li 2011) using the option “consensus”. Outgroup sequences for *Hemiprocne comata* and *Aegotheles bennettii* were obtained from the B10K avian phylogenomics dataset (Feng *et al*. 2020) and added to the exon alignments along with those for *Chaetura pelagica*. Individual exon sequence alignments were aligned with MAFFT (Katoh and Standley 2013) using the option “L-INS-i” and concatenated into a final alignment with a length of 46,416 bp after filtering with trimAl, option “-gt 0.8” (Capella-Gutiérrez *et al*. 2009).

We then obtained a time-scaled tree using Bayesian time-tree estimation methods in Beast 2 version 2.6.7 (Drummond et al. 2006; Bouckaert et al. 2019). We implemented a GTR+GAMMA model of nucleotide substitution with 6 rate categories and an uncorrelated lognormal molecular clock model of rate variation across branches with 10 rate categories. We used default and minimally informative priors for most parameters including a uniform prior for the clock rate (0 - ∞). We used a Birth-Death-Sampling model for the tree prior using the Beast package BDSKY (Stadler et al. 2013), with a uniform prior for the speciation rate (0 - 100), extinction fixed at 0, and a beta distribution (11, 79) for the sampling proportion which produces a density with a mean of 0.12 that represents the proportion of the species of hummingbirds sampled in our dataset. We fixed the tree topology to that found in the main phylogenetic analysis.

We applied a calibration prior to the root of the tree, the most recent common ancestor of Trochilidae and Aegothelidae, i.e. the clade Daedalornithes (Sangster 2005). We used the fossil record of Daedalornithes to generate an empirical calibration density using the R package CladeDate (Claramunt 2022). The clade has a notable fossil record, with the earliest taxon being *Eocypselus vincenti* Harrison 1984, represented by one nearly complete skeleton and two incomplete postcranial skeletons from the Fur Formation at Isle-of-Mors, Denmark (Dyke et al. 2004, Mayr 2010), which is between 55.8 and 54.6 Ma old (Stokke et al. 2020a, 2020b). Phylogenetic analyses place *Eocypselus* at either the stem of Apodiformes or sister to Aegothelidae (Ksepka et al. 2013, Chen et al. 2019), either way, *Eocypselus* represents the older fossil of crown Daedalornithes. We compiled the first fossil occurrence of Daedalornithes in each continent to obtain a sample of occurrences (Table S7) that are geographically independent (Claramunt & Cracraft 2015) and uniformly distributed through time (Kolmogorov-Smirnov test: D = 0.40, p-value = 0.16). We then used the seven first occurrences with the Strauss-Sadler model and Monte Carlo sampling to generate a distribution of possible ages for Daedalornithes using the function clade.date (Claramunt 2022). The function then fits standard probability densities to the empirical distribution: the best-fitting function was a lognormal (mean-log = 1.7837, sd-log = 1.0572; Figure S11), which we used as the calibration prior in BEAST.

We ran two MCMC chains for 10 million generations, sampling every 5,000 generations. We evaluated stationarity by examining trace plots, convergence by comparing the two runs, and sampling sufficiency by evaluating Effective Sample Size statistics for the likelihood and other relevant parameters in Tracer 1.7.1 (Rambaut et al. 2018). Finally, we combined the results of the two runs after discarding the first 10% as burn in and obtained the maximum clade credibility tree and node age summary statistics in Tree Annotator (Bouckaert et al. 2019).

## Results

### Target enrichment

More than 70% of the target regions were covered in all 46 captured species, and more than 90% in 38 species (Figures S3 and S4). The individual *de novo* assemblies ranged in size from ∼20 Mb to ∼100 Mb (Table S6). ∼8 Mb could be mapped with LASTZ to the Anna’s hummingbird (*Calypte anna*) reference genome (Zhang et al. 2014) resulting in matches with 94-99% identity (Tables S3, S4, and S5).

### Phylogenetic reconstruction

The major phylogenetic subdivisions within hummingbirds identified more than 20 years ago with DNA-DNA hybridization studies (Bleiweiss et al. 1994) were confirmed in our mtDNA genome and nuDNA gene trees using both a concatenation approach with RAxML (Stamatakis 2014) (Figure 1) and the multi-species coalescent approach implemented in ASTRAL (Mirarab et al. 2014) and ASTRAL-II (Mirarab and Warnow 2015) (Figure S5). All phylogenetic trees recovered the main subgroups of hummingbirds with 100% bootstrap values (using the chimney swift, *Chaetura pelagica* as outgroup). Furthermore, all trees have identical topologies for Topazes, Hermits, Mangoes, Bees, and Emeralds. Mangoes are sister to Brilliants, Coquettes, Emeralds, the genus *Patagona*, Mountain Gems, and Bees, and the two latter are sister clades in all phylogenies. Moreover, the phylogenies built using subsets of the nuDNA genes (Table S3) present the same topology between the main groups of hummingbirds with high confidence (Figure 1 and Figures S5 and S6). The three subsets correspond to: (i) the 2949 genes that were successfully captured in a minimum of 8 species, (ii) the 1987 genes for which we could assign a high confidence chimney swift ortholog as determined in (Jarvis et al. 2014), and (iii) the 741 genes within the latter that produced trees with average support above 50%. Despite consistencies in species trees estimated using various methods and datasets, there are significant conflicts among nuDNA gene trees (Figure S7), and between gene trees and the species tree (Figures 2, S1 and S2). The GC content (Figure S8) was quite stable across the three codon positions and making it less likely that codon biases resulted in incorrectly estimated gene or species trees, which is further confirmed by the topology obtained using RY-coding (Figure S9) is overall in agreement with the other nuDNA species trees. In the phylogenetic tree constructed from complete mtDNA genomes (Figure 3), the topological relationships between and within the main clades were recovered with high bootstrap confidence, except for the Brilliants and the placement of the genus *Patagona*. However, we observed a mito-nuclear discordance relative to the placement of Coquettes and Brilliants. Both our complete mtDNA genome tree and the tree of McGuire et al. (McGuire et al. 2014), which was based on 6,461 bp corresponding to concatenated nuDNA and mtDNA genes plus flanking tRNAs, posit Coquettes and Brilliants as sister clades. Accordingly, a phylogeny built using the genes that can be assigned to the Z sex chromosome places the Brilliants as sister to Coquettes (Figure S10). Yet, in our nuDNA gene phylogenies using either concatenation (Stamatakis 2014) or ASTRAL (Mirarab et al. 2014), Coquettes are sister to the clade containing Brilliants, the genus *Patagona*, Bees, Emeralds, and Mountain Gems (Figure S5). The Giant Hummingbird complex is again confirmed to be the sister to the large clade composed of Bees, Emeralds and Mountain Gems. Our mtDNA phylogeny places the genus *Patagona* as sister to a clade formed by Bees and Mountain Gems, albeit with a low bootstrap value of 68%. There are also some noteworthy differences in the within-group topology. Within Bees, Archilochus is sister to a clade containing *Calypte* and *Selasphorus* in both ASTRAL and concatenation nuDNA gene trees, whereas the mtDNA tree presents a polytomy with an unresolved placement of *Archilochus*. Within Mountain Gems, *Heliomaster* is sister to the clade containing *Lampornis* and *Panterpe* in all nuDNA topologies with 100% bootstrap support, except for the one corresponding to the smallest gene subset (Figure S5 and Figure S6), and the support is 90% in the mtDNA tree. *Lampornis* was previously considered as sister to the clade containing *Heliomaster* and *Panterpe* (McGuire et al. 2014), which is also what we find in the topology built with 741 concatenated genes. Within the Coquettes, our concatenation nuclear-based trees identify *Phlogophilus* and *Heliangelus* as sister clades, as do the ASTRAL (Mirarab et al. 2014) trees built using the subsets containing 741 and the 1987 genes (Figure S5). McGuire et al. (McGuire et al. 2014) identified *Phlogophilus* as the sister to a larger clade containing *Heliangelus* and 13 additional genera. We see a similar scenario in the mtDNA tree, where *Phlogophilus* is sister to *Lophornis*, the latter being basal to all Coquettes in the other trees. Within Brilliants, our nuDNA-based trees identify *Heliodoxa* as the sister to the clade that contains *Ocreatus, Boissonneaua, Aglaeactis, Coeligena*, and *Ensifera*. In the McGuire et al. (McGuire et al. 2014) tree, *Aglaeactis* is sister to the clade containing *Ocreatus, Boissonneaua, Heliodoxa, Coeligena*, and *Ensifera*. In our mtDNA tree, *Coeligena* is sister to *Ocreatus, Boissonneaua, Heliodoxa, Aglaeactis*, and *Ensifera*. There is considerable discordance between independent nuDNA gene trees within Brilliants (Figure S1 and S2). Bootstrap values are also low within this group in the mtDNA tree. The only species that has a stable position is *Eriocnemis alinae*, as the sister of all the other Brilliants present in our data set.

**Figure 1.**
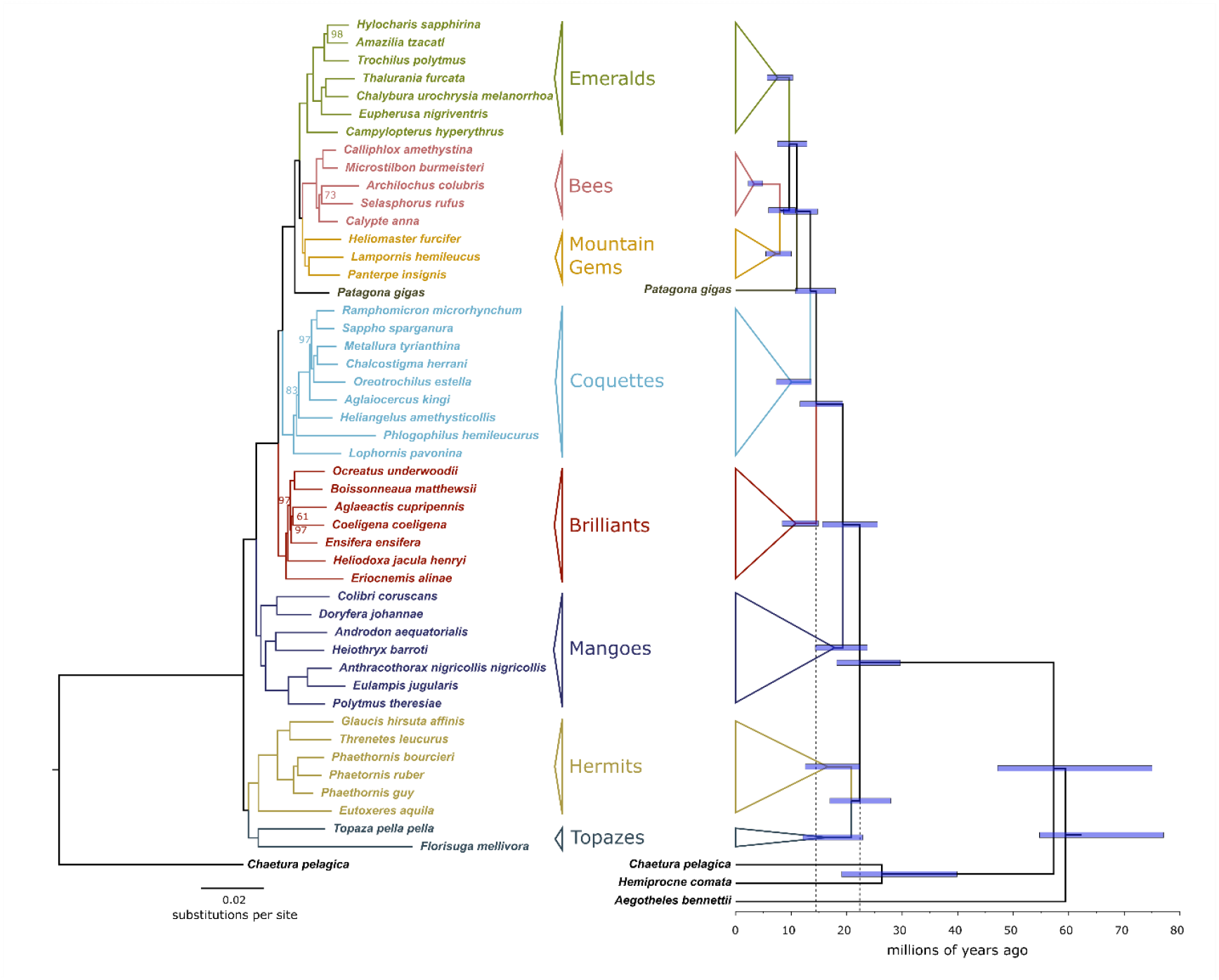
Left) Maximum-likelihood trees inferred by RAxML using concatenated alignments with the three codon positions for 1987 genes corresponding to an average of 2.9 Mb alignment size Bootstrap values below 100 are shown on branches. Right) Time-calibrated tree inferred in BEAST 2; node bars indicate 95% highest posterior density intervals for divergence time estimates.

**Figure 2.**
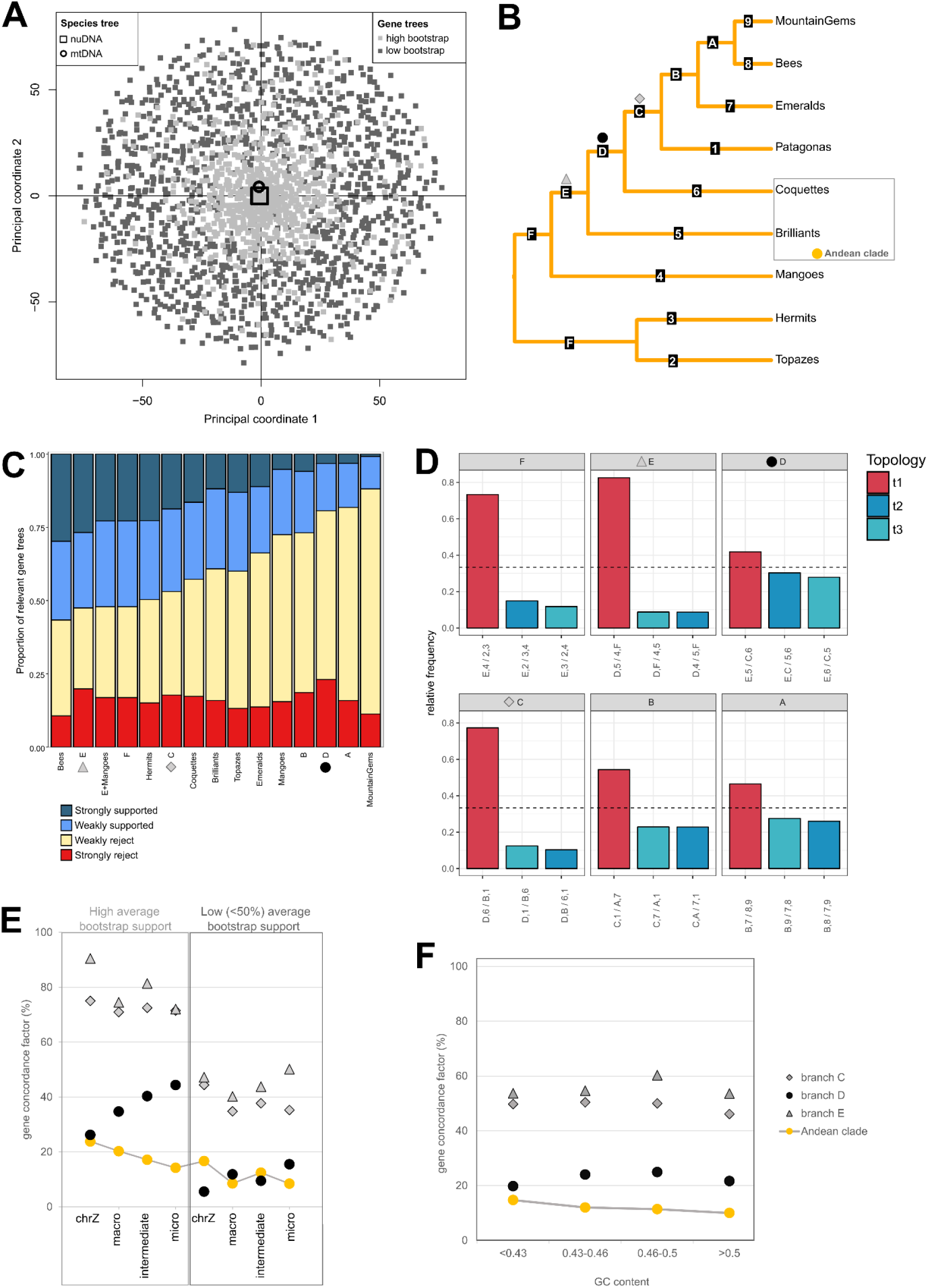
A) MDS plot reflecting the gene tree distance determined in TreeCl, highlighting those with high or low (<50%) average bootstrap support value. B) The ASTRAL tree with major clades collapsed and labels on each depicted branch. C) The graph shows the percentage of genes that support or reject each branch weakly (<75% bootstrap support) or strongly (>=75%). D) Gene tree incongruence statistics. The bargraph shows gene tree quartet frequencies for alternative branching orders for six main branches of the ASTRAL species tree estimated on all 2949 gene trees with the third codon position included (middle). Red bars represent the ASTRAL topology; blue bars represent the frequencies of alternative branching orders. Dashed horizontal lines mark equal quartet frequencies at 1/3. Each panel is labeled by a branch and each alternative topology on the bar graph is labeled according to the tree in the middle; for example, the alternative bar labeled “E,2 / 3,4” under panel F puts Hermits and Mangoes together and moves Topazes as sister to the remaining hummingbirds. E) The concordance factor was calculated by grouping gene trees by chromosome category. F) The concordance factor is shown for gene trees binned by GC content.

**Figure 3.**
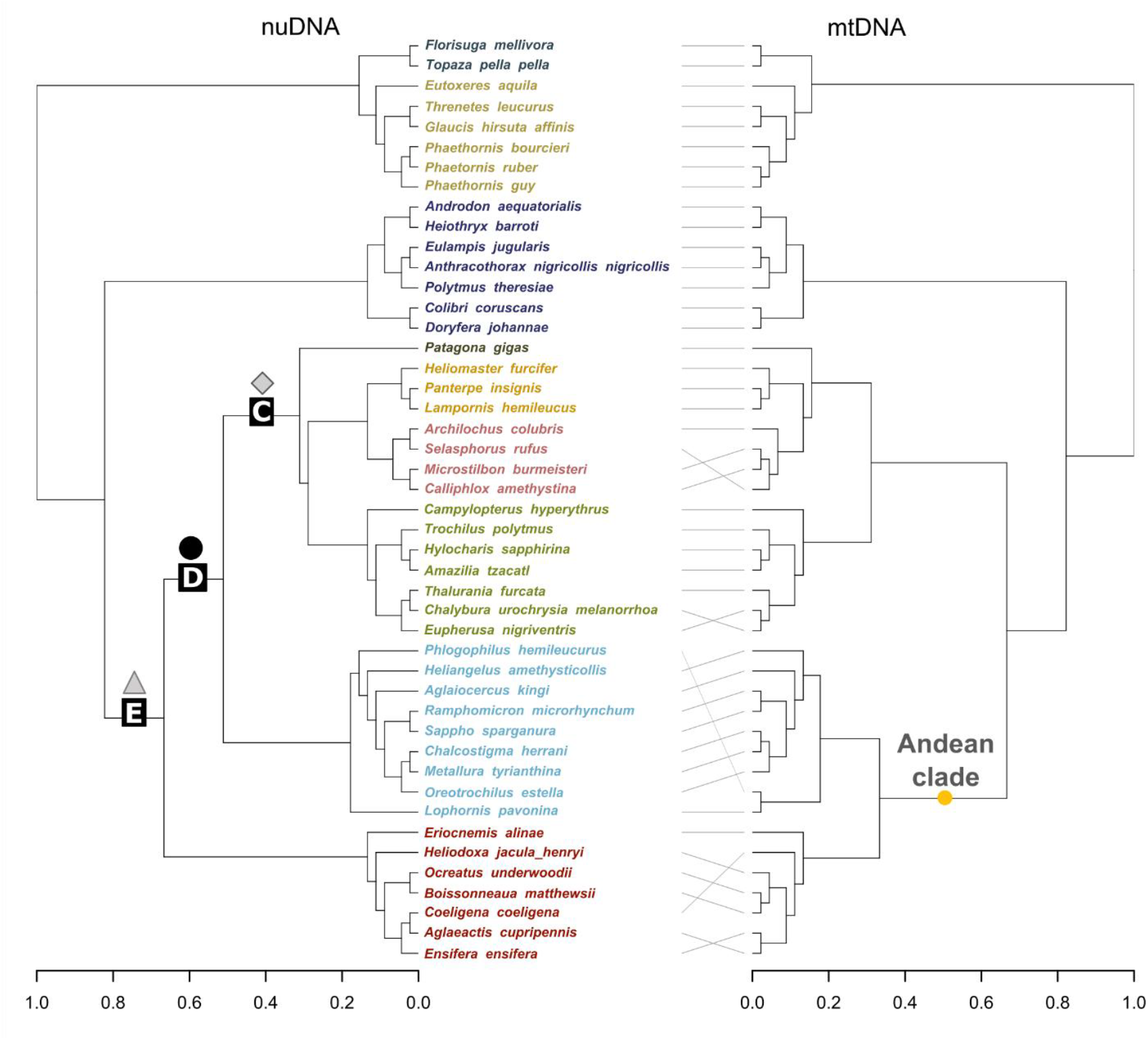
Tanglegram highlighting the topological correspondence between the nuDNA and mtDNA phylogenies (branch labelling as in Figure 2).

### Phylogenetic discordance

Gene-tree ordination based on geodesic distances revealed a single, centrally concentrated cluster of gene trees in MDS space, with no separation associated with bootstrap support (Figure 2A). The nuDNA species-tree topology occupied a central position within this cluster, closely aligned with the dominant multilocus signal, whereas the mtDNA topology was slightly offset from the centroid, indicating modest topological divergence. Despite this displacement, the absence of distinct clusters suggests that gene-tree heterogeneity is continuous rather than structured into discrete alternative topologies. The concordance between the nuDNA topology and the locus-based signal is higher for gene trees with high average bootstrap support (Figures 2E and 2F), and this is consistent across chromosomal assignment and GC content binning.

### Molecular dating

Our dating analyses, based on a reduced but topologically consistent 46,416 bp dataset and refined through an ML approach, indicate that modern hummingbirds originated ∼22 Ma and that the major Andean radiation occurred from ∼16 Ma to ∼14 Ma— a timeframe supported by broad HPD intervals—while also revealing younger divergence ages across several clades (Figure 1) when compared to the McGuire et al. tree (McGuire et al. 2014).

## Discussion

An analysis of 2949 nuDNA loci and complete mtDNA genomes confirmed the monophyly of the nine major clades of hummingbirds described in previous studies (Bleiweiss 1998b; McGuire et al. 2014). There is, however, one clear difference in the timing of diversification of Coquettes and Brilliants when we compare the nuDNA and mtDNA trees. The nuDNA tree suggests that Brilliants and Coquettes are sequentially derived from lowland ancestors, as indicated by Bleiweiss’s phylogeny (Bleiweiss 1998b), whereas the mtDNA suggests that Brilliants and Coquettes are sister taxa as in the McGuire et al. tree (McGuire et al. 2014). The recovery of the same overall topology with high bootstrap values for most nodes using different subsets of the nuDNA markers, and the use of full mtDNA genome information allows us to confidently state that there is a true discordance between the nuDNA vs mtDNA and Z-linked markers. Discordance between differentially inherited markers can simply result from stochastic patterns of lineage sorting, but it can also be indicative of introgression (Lavretsky et al. 2014). Many suspected intra- and inter-generic hybrids among Andean Brilliants and Coquettes are known (Graves 1990, 1996, 1997, 1998, 1999, 2001, 2004; Graves and Zusi 1990), but only a single hybrid zone is known in the Trochilidae, between streamertail hummingbirds (*Trochilus*) in Jamaica (Graves 2015, Judy et al. 2025). The process that led to the observed discordance between nuDNA and mtDNA markers regarding the relationship between Coquettes and Brilliants must have occurred early during the divergence of the two groups, both of which are monophyletic in both nuDNA and mtDNA trees. The very short branch leading to the Andean clade in the mtDNA tree could reflect a rapid burst of early diversification. In this scenario, incomplete lineage sorting of the mitochondria could explain the mito-nuclear discordance. Note that in the ASTRAL tree, several branches had very short lengths in coalescent units (Figure S6), e.g. the branch separating Brilliants and other crown species had a length of only 0.14 in coalescent units. Furthermore, the percentage of gene tree quartets supporting that branch is 42%, with the other two alternative hypotheses possible for an unrooted quartet present in 30% and 28% of the genes (Figures 2D and S2). Alternatively, the observed mito-nuclear discordance might have resulted from introgression between the ancestors of Coquettes and the incipient Brilliant species. Mito-nuclear discordance within an *Amazilia* species complex has been linked to introgression occurring in the edges of the species ranges (Jiménez and Ornelas 2015). It is worth noting that Coquettes and Brilliants represent ∼30% of extant hummingbird taxa and most have Andean distributions. Moreover, Coquettes and Brilliants are the prevalent hummingbird groups above 3,000 m (Graham et al. 2009), having successfully colonized and adapted to some of the most challenging environments in the Western Hemisphere (Altshuler et al. 2004; Projecto-Garcia et al. 2013; Stiles 2008). The rise of the Andes created vast unexploited habitats that provided the perfect setting for divergence between Coquettes and Brilliants. Nevertheless, the quartet frequencies (Figure 2D and Figure S2) are consistent with a scenario in which incomplete lineage sorting alone accounts for the observed gene-tree discordance, with gene-tree heterogeneity arising from coalescent stochasticity associated with short internodes, without invoking introgression or other sources of non-tree-like evolution.

Our divergence time estimates agree closely with some previous estimates. For example, both our new estimates and those of McGuire et al. (2014) place the most recent common ancestor of extant hummingbirds at 22 Ma. This is remarkable given the different sources for the time calibration in these two studies, one based on rates of molecular evolution estimated for Hawaiian honeycreepers (McGuire et al. 2014), and the other based on the fossil record of Apodiformes and relatives (this study). Furthermore, our dating analyses reveal that hummingbird diversification follows a stepwise Andean pattern, with a major radiation occurring around ∼14 Ma. This time coincides with a phase of substantial Andean uplift resulting in the rise of the proto– Cordillera Oriental and Altiplano to roughly half their modern elevation by the Middle Miocene and continued rapidly over the last ∼12 million years (Graham 2009), which fits our timing of the diversification of Emeralds, Bees and Mountain Gems (Figure 1). This uplift reshaped regional atmospheric circulation and rainfall regimes, producing a patchwork of cool, mist-laden cloud-forest microhabitats along the rising Andean slopes (Fjeldså et al. 2012). In addition to Andean uplift, this period coincides with the Middle Miocene Climatic Transition, a period of global cooling that produced the expansion of open and cold habitats and around the world (Flower & Kennett 1994), and rapid diversification of plant clades that may have provided nectar resources (Dellinger et al. 2024, Barreto et al. 2024). The associated burst of plant diversification—particularly among lineages evolving tubular, nectar-rich flowers—opened new ecological opportunities for hummingbirds to occupy emerging elevational niches, often displacing insects as the primary pollinators in these cool, humid environments (Sonne 2019; McGuire et al. 2014, Lagomarsino et al. 2016, Barreto et al. 2024). Together, these processes likely accelerated the tempo of hummingbird speciation by coupling geological uplift with the development of highly structured floral communities and increasingly fragmented montane habitats (Graham 2009; Fjeldså et al. 2012).

## Conclusions

Here we present hummingbird phylogenies using nuDNA and complete mtDNA genomes, both recovering the monophyly of hummingbirds and its main subgroups with 100% bootstrap values. However, there are differences in the topology of trees built using autosomal genes when compared to the sex chromosome Z and the mtDNA, namely the relative position of Coquettes and Brilliants. The short branch separating Brilliants and the clade containing Coquettes, Emeralds, the genus *Patagona*, Mountain Gems, and Bees suggests that the early diversification was very rapid, and we show that incomplete lineage sorting contributes to the discordance, which could also explain the high discordance between gene trees relative to the same branch.

## Data availability

Data available upon request.

## Author contributions

R.R.F., G.R.G., A.A., T.W., J.F., M.T.P.G. and C.R. designed the study; G.R.G. provided the samples. I.W. performed the laboratory work. R.R.F., S.M., S.C., M.M.F, J.A.S.C., C.L., and F.P. analyzed the data, and R.R.F. wrote the manuscript with contributions from all the authors.

## Acknowledgments

We thank the National Museum of Natural History, Smithsonian Institution (USNM) and the Louisiana State University Museum of Natural Sciences (LSUMZ) for providing samples. We thank Ricardo Pereira and Angela M. Ribeiro for stimulating discussion and comments on the manuscript. We thank Jesper Sonne for facilitating the access to previous hummingbird phylogenies and clade annotations. Letty Salinas provided information on specimens in the Museo de Historia Natural, Lima, Peru.

