## Supplementary material for "A time-calibrated phylogeny of hummingbirds supports stepwise diversification in the Andes": Supp

### Supplementary Information

Supplementary Methods

Supplementary Figures S1 to S11

Supplementary Tables S1 to S7

### Supplementary Methods

#### Bait design

The set of 2949 genes was selected according the following criteria:

- The genes corresponded to one-to-one orthology between chicken and finch as annotated in ENSEMBL version 66 annotation;
- Presence of a functional annotation (GO assignment);
- Annotated as “known” genes;
- Between 50-95% similarity chicken-finch;
- >85% similarity hummingbird-swift;
- Discarded hummingbird gene sequences that have undetermined bases (Ns).
- The probes were design according to the following:
- 100 bp of intron flanking sequence was added to each exon;
- Overlapping bases were merged;
- The humming bird scaffolds were masked for low complex regions and repetitive sequences of the chicken and probes that had 50% of their sequences masked were discarded;
- The probes have an average length of 120 bps with a tiling of 40 bps;
- Probes smaller than 80 bp were discarded.

The final set contained 166322 probes corresponding to 2949 genes and summing up to approximately 20 Mb.

#### DNA extraction and capture-enrichment

Muscle biopsy samples were stored in RNAlater at -20°C until DNA was extracted. Extractions were performed with the DNeasy Blood & Tissue kit from Qiagen according to the manufacturer’s instructions. In the final step each sample was eluted in 100µl of elution buffer. DNA integrity was evaluated by gel electrophoresis (5µl sample, 1% agarose, 100V, 30min, 1kb ladder), and concentration was determined with the Qubit® 1.0 Fluorometer from Invitrogen using 1µl of sample and following the user manual. Once sufficient quality was determined, the DNA extracts were sonicated at the HIGH setting for 11 cycles of 30/30 seconds on the Bioruptor® Ultrasonicator from Diagenode. This was sufficient to fragment the extracted genomic DNA to an average length of 200bp, which was measured with the Agilent High Sensitivity DNA Kit for the 2100 Bioanalyzer from Agilent Technologies. 50µl of each sample was used to build NGS libraries using the NEBNext DNA Sample Prep Master Mix Set 2 from New England Biolabs and Illumina specific adapters. Enrichment PCRs of each library were performed using unique sets of indexing primers and Platinum® Taq DNA Polymerase High Fidelity from Invitrogen (50µl reactions with the following concentrations: 1x High Fidelity PCR Buffer, 0.2mM dNTPs, 2mM Magnesium Sulphate, 0.4µM forward primer, 0.4µM indexed reverse primer, ddH20 and 2U polymerase per reaction) to reach a minimum of 350ng of DNA for the following capture protocol. Library DNA concentrations were also measured on the Bioanalyzer. The PCR amplified libraries were then concentrated down to a volume of 3.5µl, in preparation for the capture protocol, by using a vacuum centrifuge. The custom RNA baits were synthesized by MYcroarray and used with the MYbaits custom target enrichment kit from the same company for the solution-based capture according to manufacturer instructions (MYbaits protocol version 1.3.6 - 06/01/2012). Following capture the enriched libraries were amplified, using the same primers and conditions as previously described with the Platinum® Taq kit from Invitrogen, until concentrations were sufficient to combine in equimolar balanced pools before sequencing.

#### Supplementary Figures

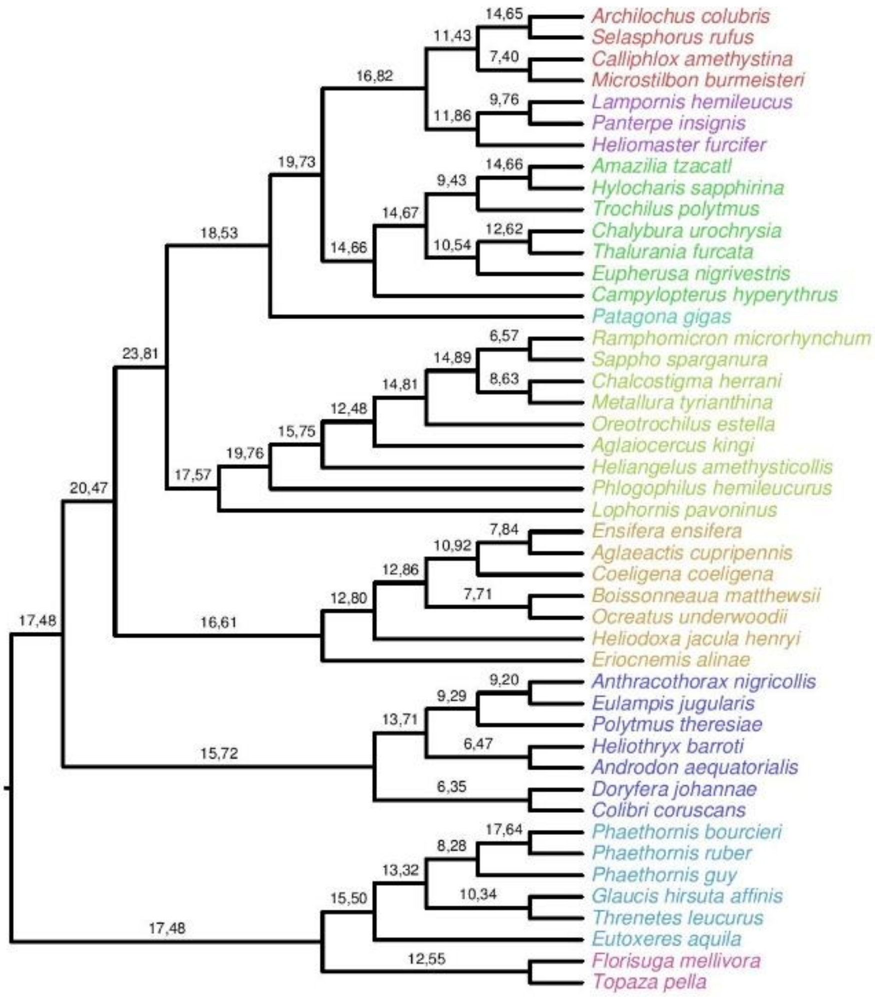


**Figure S1.** Comparison between gene trees and the species tree with the full dataset (2949 genes). The first number gives the percentage of genes that conflicted with a species tree branch at 75% support. The second number is the percentage of genes that had conflict with a branch, whether that conflict was highly supported or not. There is a lot of unsupported gene tree conflict, and there is also a fair bit of supported gene tree conflict (10-20% typically).

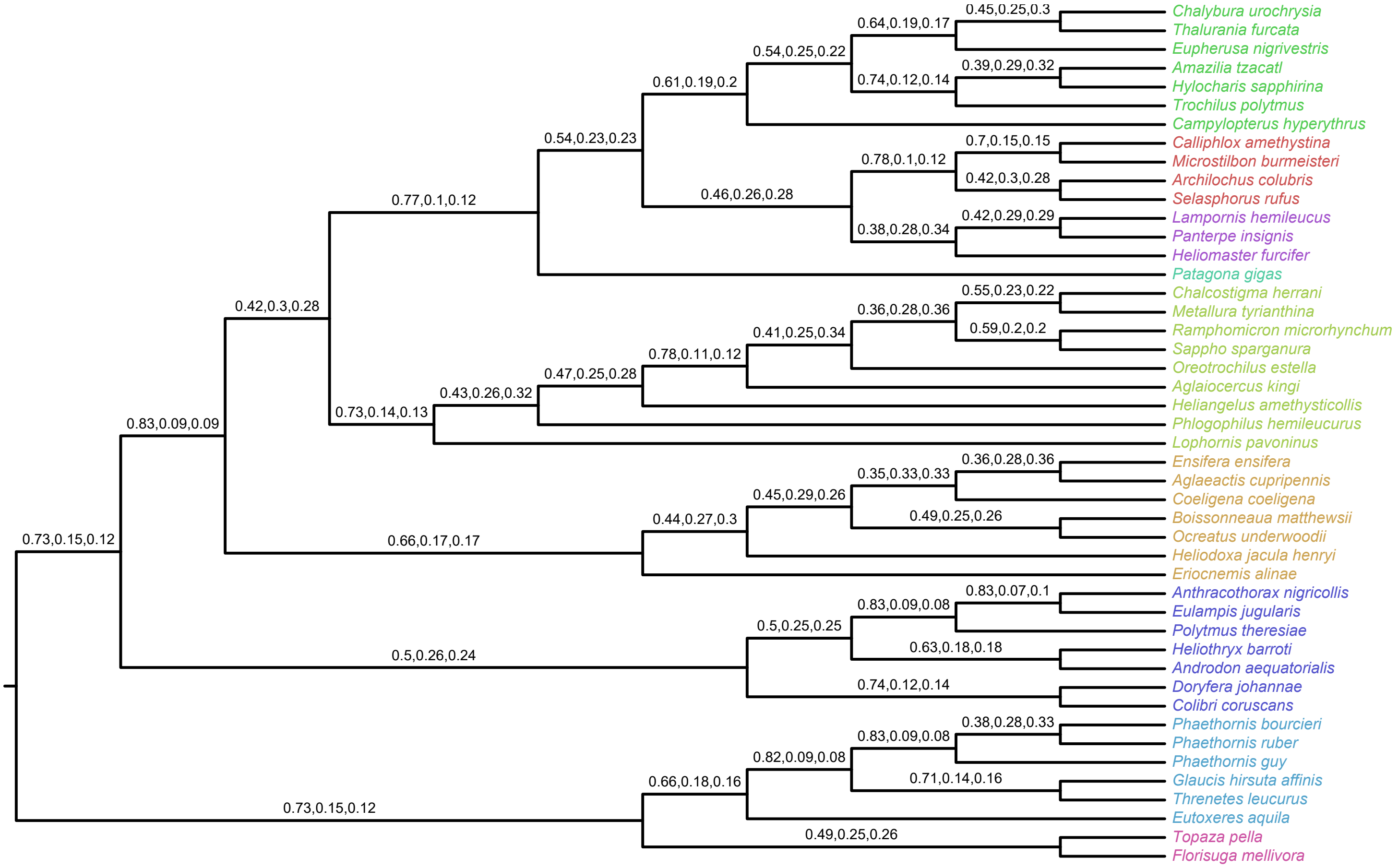


**Figure S2**. Quantifying gene tree incongruence using quartet support values. For each branch in the ASTRAL tree, we show the percentage of gene tree quartets around that branch that are compatible with the ASTRAL topology (the first number) or the two alternative hypothesis possible for an unrooted quartet (second and third numbers).

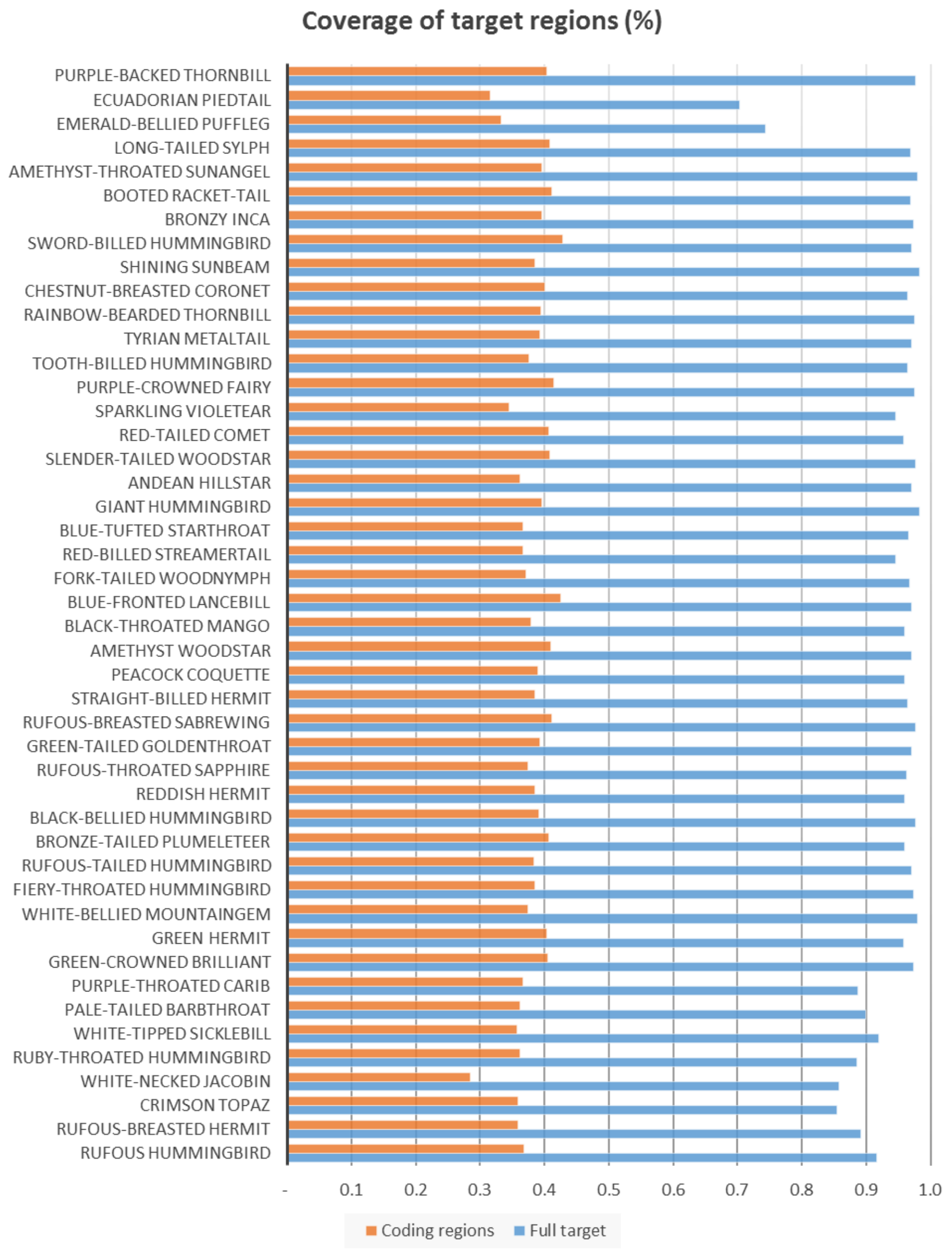


**Figure S3.** Sequence coverage of the target regions designed using Anna's hummingbird gene sequences. The target regions contained exons and flanking regions (UTRs and introns). The blue bars indicate the percentage of the total length of the targets regions that was covered per species. The orange bars indicate the percentage of the total length of the targets regions that was covered per species that corresponds to coding regions.

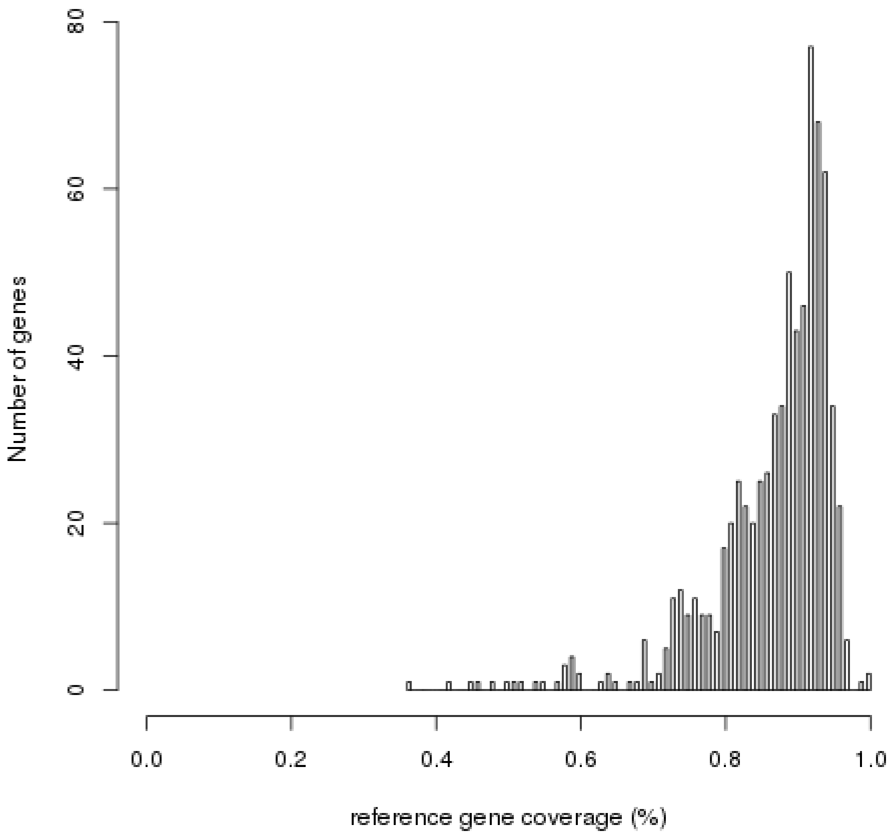


**Figure S4.** Distribution of the coverage of the target Anna's hummingbird genes. For most genes more than 80% of the coding region was successfully captured.

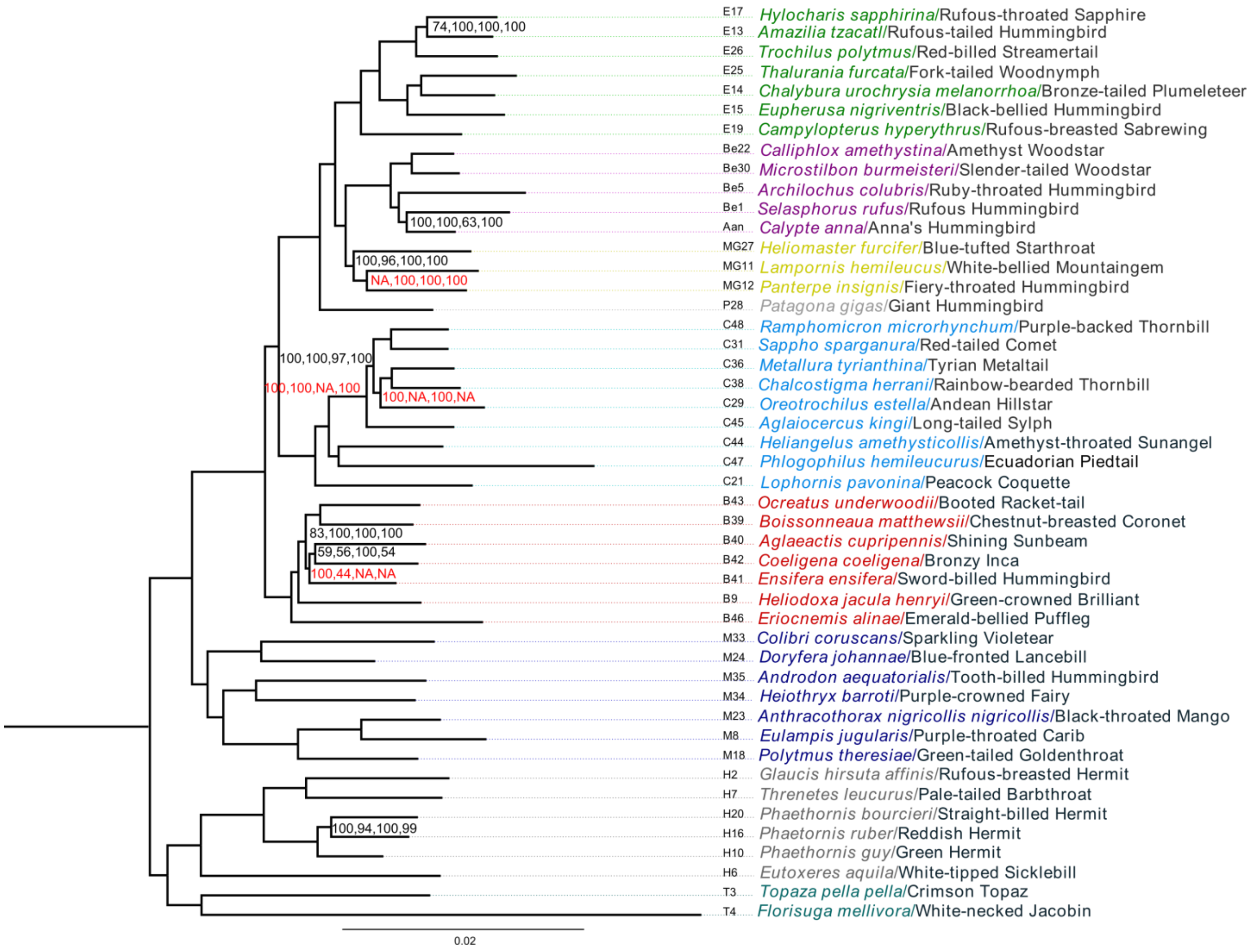


**Figure S5.** Consensus topology obtained for 4 trees using the 2 datasets that include high confidence swift orthologs. Trees were obtained with 741 genes or 1987 genes using either a concatenation approach with RAxML or the multi-species coalescent approach implemented in ASTRAL. The tree is rooted with the chimney swift, which is not visible in the figure. The bootstrap label order for the species tree: 741g_raxml, 741g_astral, 1987g_raxml, 1987g_astral. When the bootstrap value is not 100% in all analyses, the bootstrap values are shown. The bootstrap values in red highlight the branches that are not represented in at least one of the 8 topologies (indicated by a "NA").

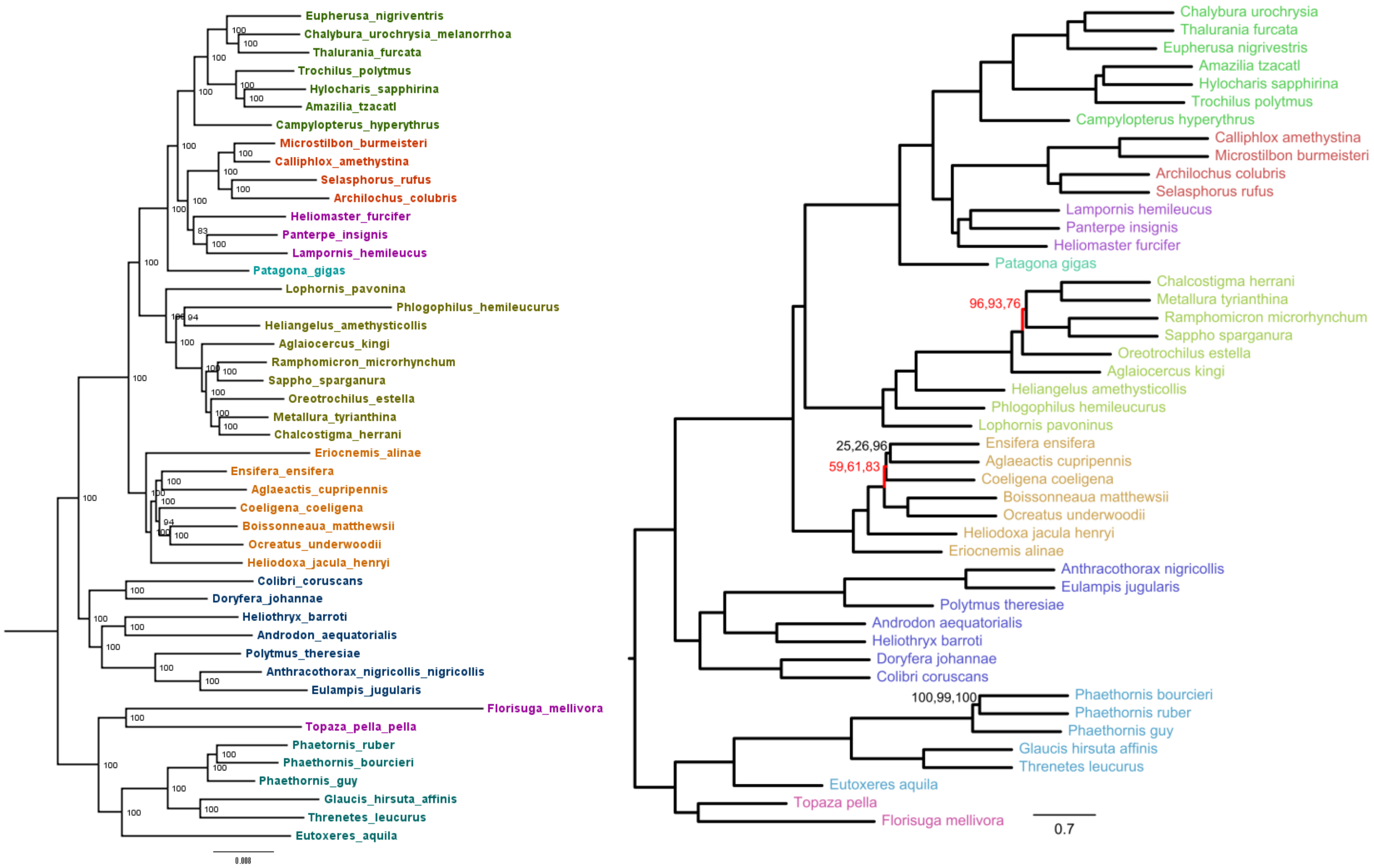


**Figure S6.** Topologies obtained with the full dataset (2949 genes), using all three codon positions and no outgroup. Left: RAxML, concatenated data set. Right: Astral tree with internal branch lengths in coalescent units (terminals shown arbitrarily). Three types of support are shown for ASTRAL: local posterior probabilities, site-only bootstrap, and gene+site bootstrap; all three values are 100% for branches with no designation and are shown in that order for the remaining branches. The two branches that differed with concatenation are shown in red.

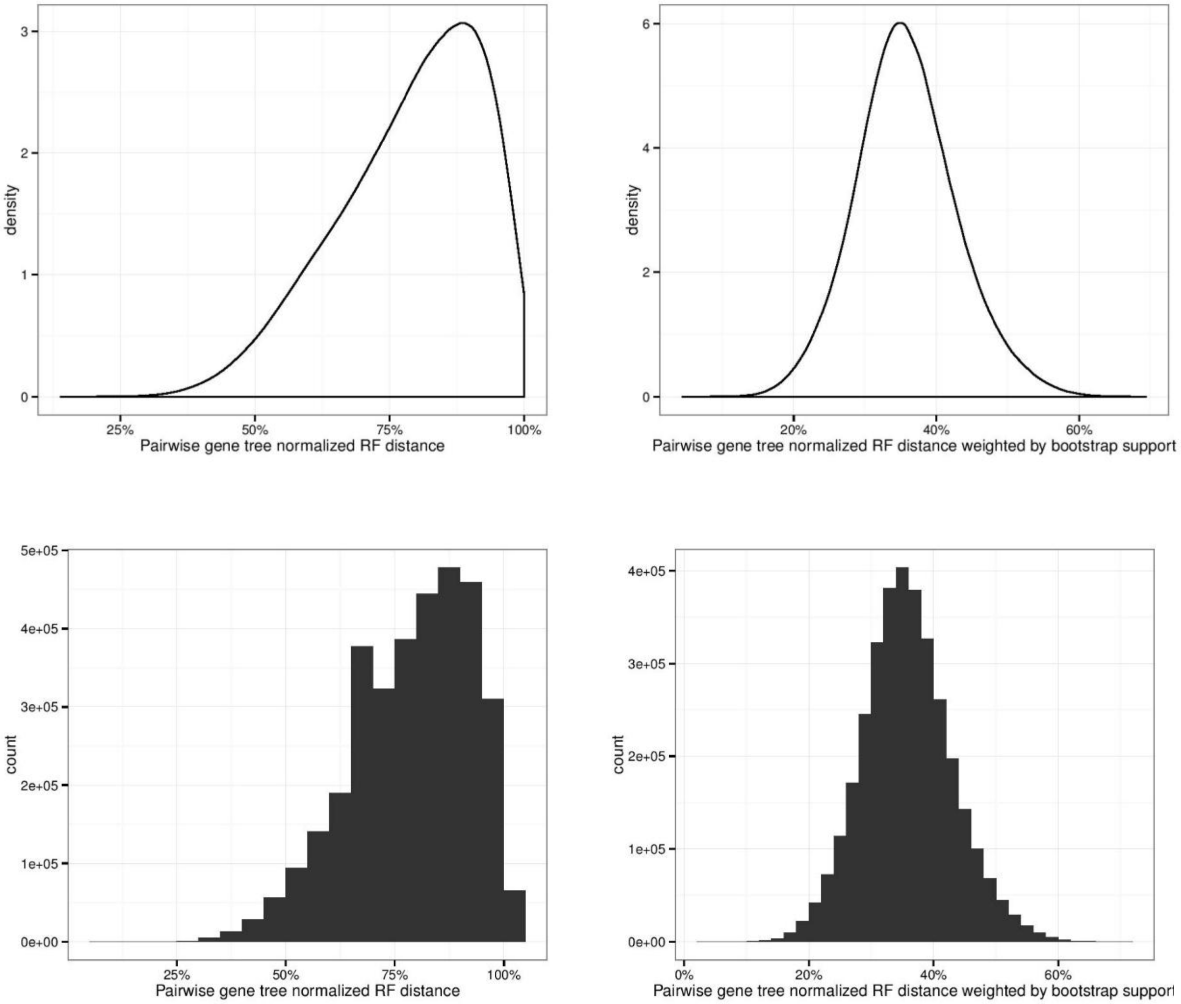


**Figure S7.** Pairwise gene tree distances using with the full dataset (2949 genes). There is a lot of discordance between gene trees (typically between 60% to 90% conflict) but once conflict is weighted by bootstrap support (the weighting schema implemented in RAxML; plots on the left) it is reduced to 20% to 60%. 1085 genes have >50% support, and the average of averages is around 45%.

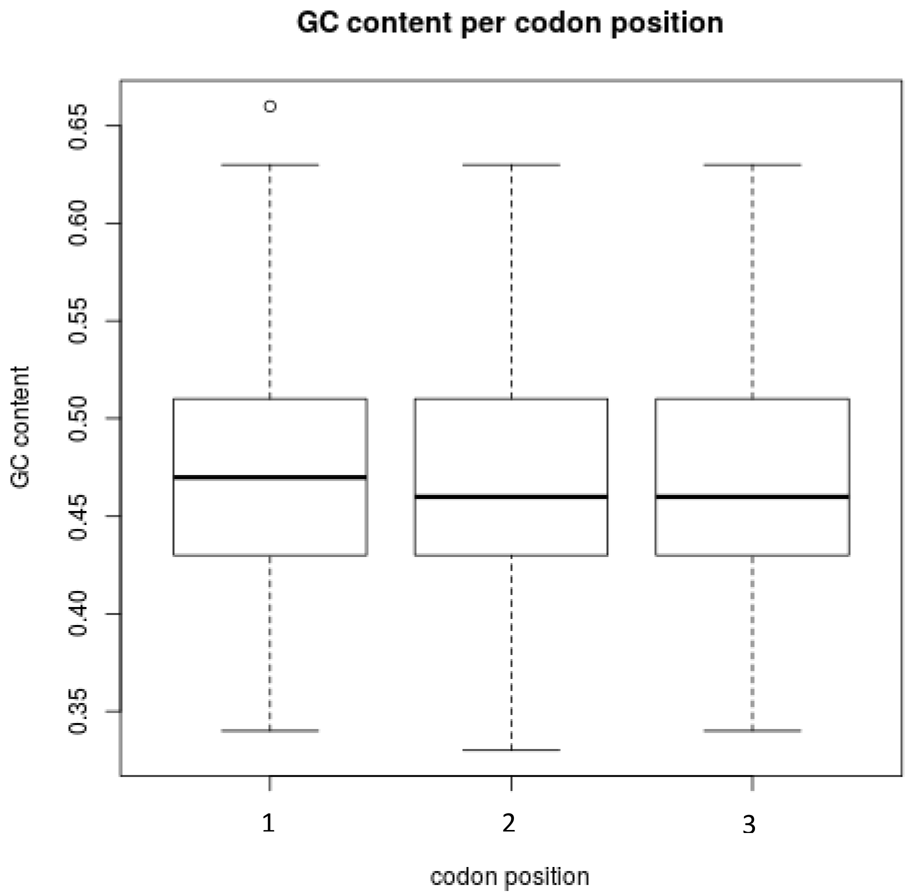


**Figure S8.** GC content distribution per gene, position and species in the 741 gene dataset.

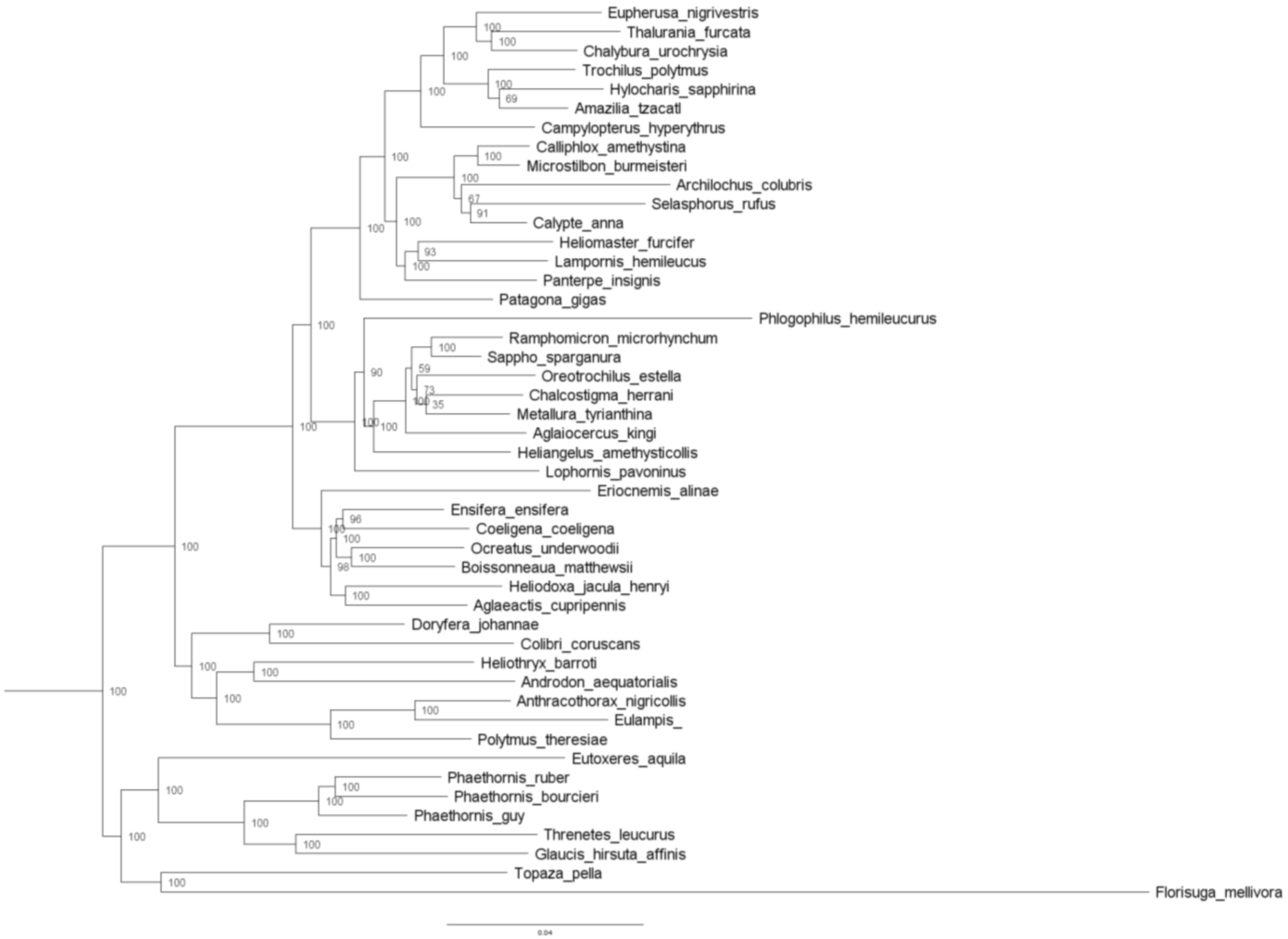


**Figure S9.** Topology obtained using RY-coding with the concatenated sequences of 741 genes in RAxML. The tree is rooted with the chimney swift, which is not visible in the figure. The node labels correspond to the bootstrap value.

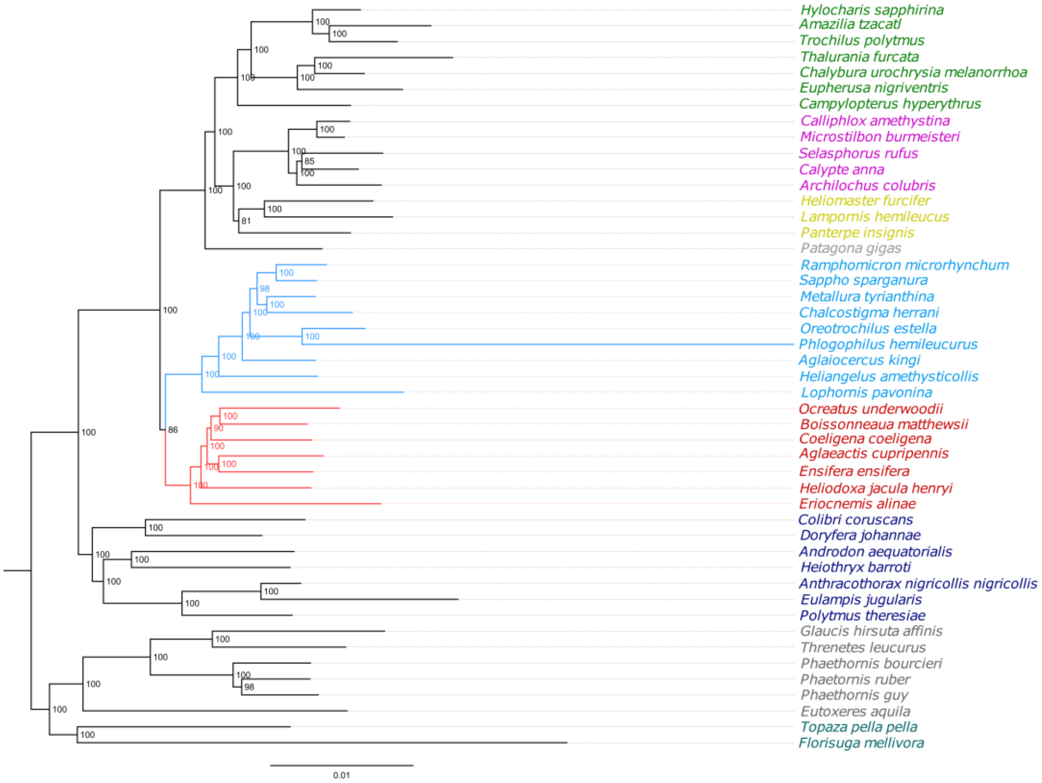


**Figure S10.** Topology obtained using the genes assigned to the sex chromosome Z (by orthology to the chicken). The trees were determined in RAxML using all 3 codon positions. The tree is rooted with the swift, which is not visible in the figure.

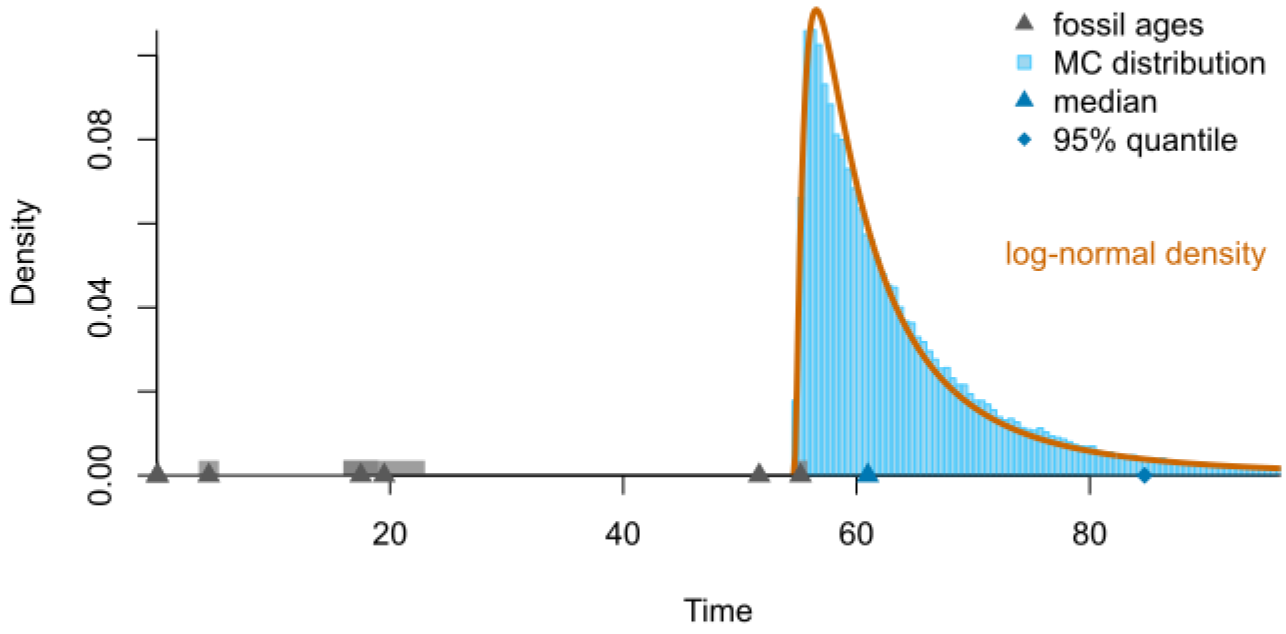


**Figure S11.** Empirical calibration density generated using CladeDate (Claramunt 2022). First temporal occurrence of Daedalornithes in each continent are show with black triangles, with grey bars represent the entire time intervals (from Table S7). Sky-blue histogram represents the distribution of ages generated by clade.date. Blue symbols are quantiles of this distribution, and the red line is a lognormal probability density distribution fit to the Monte Carlo distribution, which was used as calibration prior in BEAST2.

#### Supplementary Tables

**Table S1.** Museum identification of the samples used in this study. Tissues were obtained from the National Museum of Natural History, Smithsonian Institution (USNM)(n = 33 species) and the Louisiana State University Museum of Natural Sciences (LSUMZ)(n = 13 species). Voucher specimens for some of the LSUMZ tissue samples are deposited in the Academy of Natural Sciences of Drexel University (ANSP) and the Museo de Historia Natural (MUSM), Lima, Peru. Some MUSM specimens on loan to LSUMZ are designated by field numbers. Sexual determination was made in the field (F: female, M: male, U: unknown).

| Exemplar species | Sex | Tissue number | Museum Catalog Number | Collection site (country: department, state, or province) |
|---|---|---|---|---|
| *Aglaeactis cupripennis* | F | B32232 | LSUMZ 169641 | Peru: Cajamarca |
| *Aglaiocercus kingi* | M | B33579 | RCF 1108 (field number) | Peru: Cajamarca |
| *Amazilia tzacatl* | M | B01184 | USNM 607716 | Panama: Bocas del Toro |
| *Androdon aequatorialis* | U | B12025 | ANSP 180239 | Ecuador, Esmeraldas |
| *Anthracothorax nigricollis* | F | B13779 | USNM 627476 | Guyana: Upper Takutu-Upper Essequibo |
| *Archilochus colubris* | M | B17004 | USNM 633824 | United States: Virginia |
| *Boissonneaua matthewsii* | M | B32173 | LSUMZ 169698 | Peru: Cajamarca |
| *Calliphlox amethystina* | M | B13639 | USNM 627430 | Guyana: Upper Demerara-Berbice |
| *Campylopterus hyperythrus* | M | B15813 | USNM 626797 | Guyana: Cuyuni-Mazaruni |
| *Chalcostigma herrani* | M | B31966 | RCF 608 (field number) | Peru: Cajamarca |
| *Chalybura urochrysia* | F | B03247 | USNM 608925 | Panama: Bocas del Toro |
| *Coeligena coeligena* | M | B32790 | USNM 169696 | Peru: Cajamarca |
| *Colibri coruscans* | M | B8406 | USNM 645731 | Argentina: Tucuman |
| *Doryfera johannae* | M | B19035 | USNM 632743 | Guyana: Cuyuni-Mazaruni |
| *Ensifera ensifera* | M | B32689 | CCW 268 (field number) | Peru: Cajamarca |
| *Eriocnemis alinae* | M | B44251 | MUSM 24969 | Peru: San Martin |
| *Eulampis jugularis* | F | B02105 | USNM 607198 | St. Vincent and the Grenadines: St. Vincent |
| *Eupherusa nigrivestris* | M | B01489 | USNM 613299 | Panama: Chiriqui |
| *Eutoxeres aquila* | F | B01428 | USNM 542826 | Panama: Chiriqui |
| *Florisuga mellivora* | F | B18919 | USNM 632616 | Guyana: Cuyuni-Mazaruni |
| *Glaucis hirsuta affinis* | M | B01232 | USNM 607666 | Panama: Bocas del Toro |
| *Heliangelus amethysticollis* | M | B33947 | LSUMZ 172046 | Peru: Cajamarca |
| *Heliodoxa jacula henryi* | M | B01518 | USNM 607598 | Panama: Chiriqui |
| *Heliomaster furcifer* | M | B21035 | USNM 636130 | Uruguay: Artigas |
| *Heliothryx barroti* | U | B12039 | ANSP 180243 | Ecuador, Esmeraldas |
| *Hylocharis sapphirina* | M | B11759 | USNM 625393 | Guyana: Upper Takutu-Upper Essequibo |
| *Lampornis hemileucus* | M | B01458 | USNM 607604 | Panama: Chiriqui |
| *Lophornis pavoninus* | M | B15968 | USNM 626831 | Guyana: Cuyuni-Mazaruni |
| *Metallura tyrianthina* | F | B31741 | LSUMZ 169718 | Peru: Cajamarca |
| *Microstilbon burmeisteri* | F | B8262 | USNM 645516 | Argentina: Tucuman |
| *Ocreatus underwoodii* | M | B33042 | DFL 991 (field number) | Peru: Cajamarca |
| *Oreotrochilus estella* | M | B8179 | USNM 645497 | Argentina: Tucuman |
| *Panterpe insignis* | M | B01499 | USNM 607605 | Panama: Chiriqui |
| *Patagona gigas* | M | B7984 | USNM 645459 | Argentina: Tucuman |
| *Phaethornis bourcieri* | M | B15907 | USNM 626815 | Guyana: Cuyuni-Mazaruni |
| *Phaethornis guy* | M | B01511 | USNM 607601 | Panama: Chiriqui |
| *Phaethornis ruber* | F | B10342 | USNM 625120 | Guyana: Upper Takutu-Upper Essequibo |
| *Phlogophilus hemileucurus* | F | B44649 | MUSM 24973 | Peru: San Martin |
| *Polytmus theresiae* | F | B11975 | USNM 625394 | Guyana: Upper Takutu-Upper Essequibo |
| *Ramphomicron microrhynchum* | M | B8368 | LSUMZ 128295 | Peru: Pasco |
| *Sappho sparganura* | M | B8311 | USNM 645525 | Argentina: Tucuman |
| *Selasphorus rufus* | M | B09028 | USNM 601154 | United States: Washington |
| *Thalurania furcata* | M | B18924 | USNM 632784 | Guyana: Cuyuni-Mazaruni |

| | | | | |
|---|---|---|---|---|
| *Threnetes leucurus* | M | B09213 | USNM 586316 | Guyana: Barima-Waini |
| *Topaza pella* | M | B05253 | USNM 609131 | Guyana: Cuyuni-Mazaruni |
| *Trochilus polytmus* | M | GRG 4014 | USNM 633628 | Jamaica: Portland |

**Table S2.** Capture data was obtained for 46 hummingbird species. The publicly available genomes (*) of the Anna's hummingbird (*Calypte anna*) and the chimney swift (*Chaetura pelagica*) were also used in this study.

| Exemplar species | English name | Higher order taxonomic group | Common name |
|---|---|---|---|
| ***Trochilus polytmus*** | Emeralds | Trochilini | Red-billed Streamertail |
| ***Hylocharis sapphirina*** | Emeralds | Trochilini | Rufous-throated Sapphire |
| ***Chalybura urochrysia*** | Emeralds | Trochilini | Bronze-tailed Plumeleteer |
| ***Thalurania furcata*** | Emeralds | Trochilini | Fork-tailed Woodnymph |
| ***Amazilia tzacatl*** | Emeralds | Trochilini | Rufous-tailed Hummingbird |
| ***Campylopterus hyperythrus*** | Emeralds | Trochilini | Rufous-breasted Sabrewing |
| ***Eupherusa nigrivestris*** | Emeralds | Trochilini | Black-bellied Hummingbird |
| ***Archilochus colubris*** | Bees | Mellisugini | Ruby-throated Hummingbird |
| ***Selasphorus rufus*** | Bees | Mellisugini | Rufous Hummingbird |
| ***Calliphlox amethystina*** | Bees | Mellisugini | Amethyst Woodstar |
| ***Microstilbon burmeisteri*** | Bees | Mellisugini | Slender-tailed Woodstar |
| ***Calypte anna**** | Bees | Mellisugini | Anna's Hummingbird |
| ***Heliomaster furcifer*** | Mountain Gems | Lampornithini | Blue-tufted Starthroat |
| ***Panterpe insignis*** | Mountain Gems | Lampornithini | Fiery-throated Hummingbird |
| ***Lampornis hemileucus*** | Mountain Gems | Lampornithini | White-bellied Mountaingem |
| ***Patagona gigas*** | Giant Hummingbird | Patagonini | Giant Hummingbird |
| ***Phlogophilus hemileucurus*** | Coquettes | Lesbiini | Ecuadorian Piedtail |
| ***Lophornis pavoninus*** | Coquettes | Lesbiini | Peacock Coquette |
| ***Sappho sparganura*** | Coquettes | Lesbiini | Red-tailed Comet |
| ***Oreotrochilus estella*** | Coquettes | Lesbiini | Andean Hillstar |
| ***Aglaiocercus kingi*** | Coquettes | Lesbiini | Long-tailed Sylph |
| ***Metallura tyrianthina*** | Coquettes | Lesbiini | Tyrian Metaltail |
| ***Chalcostigma herrani*** | Coquettes | Lesbiini | Rainbow-bearded Thornbill |
| ***Ramphomicron microrhynchum*** | Coquettes | Lesbiini | Purple-backed Thornbill |
| ***Heliangelus amethysticollis*** | Coquettes | Lesbiini | Amethyst-throated Sunangel |
| ***Eriocnemis alinae*** | Brilliants | Coeligenini | Emerald-bellied Puffleg |
| ***Ensifera ensifera*** | Brilliants | Coeligenini | Sword-billed Hummingbird |
| ***Boissonneaua matthewsii*** | Brilliants | Coeligenini | Chestnut-breasted Coronet |
| ***Ocreatus underwoodii*** | Brilliants | Coeligenini | Booted Racket-tail |
| ***Heliodoxa jacula henryi*** | Brilliants | Coeligenini | Green-crowned Brilliant |
| ***Coeligena coeligena*** | Brilliants | Coeligenini | Bronzy Inca |
| ***Aglaeactis cupripennis*** | Brilliants | Coeligenini | Shining Sunbeam |
| ***Eulampis jugularis*** | Mangoes | Polytmini | Purple-throated Carib |
| ***Colibri coruscans*** | Mangoes | Polytmini | Sparkling Violetear |
| ***Anthracothorax nigricollis*** | Mangoes | Polytmini | Black-throated Mango |
| ***Androdon aequatorialis*** | Mangoes | Polytmini | Tooth-billed Hummingbird |
| ***Polytmus theresiae*** | Mangoes | Polytmini | Green-tailed Goldenthroat |
| ***Doryfera johannae*** | Mangoes | Polytmini | Blue-fronted Lancebill |
| ***Heliothryx barroti*** | Mangoes | Polytmini | Purple-crowned Fairy |
| ***Glaucis hirsuta affinis*** | Hermits | Phaethornithinae | Rufous-breasted Hermit |

| Exemplar species | English name | Higher order taxonomic group | Common name |
|---|---|---|---|
| ***Threnetes leucurus*** | Hermits | Phaethornithinae | Pale-tailed Barbthroat |
| ***Eutoxeres aquila*** | Hermits | Phaethornithinae | White-tipped Sicklebill |
| ***Phaethornis guy*** | Hermits | Phaethornithinae | Green Hermit |
| ***Phaethornis ruber*** | Hermits | Phaethornithinae | Reddish Hermit |
| ***Phaethornis bourcieri*** | Hermits | Phaethornithinae | Straight-billed Hermit |
| ***Topaza pella*** | Topazes | Topazini | Crimson Topaz |
| ***Florisuga mellivora*** | Topazes | Topazini | White-necked Jacobin |
| ***Chaetura pelagica**** | | | Chimney Swift (outgroup) |

**Table S3.** Nuclear gene datasets used to build phylogenetic trees. The three subsets correspond to: *i)* the 2949 genes that were successfully captured (in a minimum of 8 species), *ii)* the 1987 genes for which we could assign a high confidence swift ortholog (used as a root), and *iii*) the 741 genes within the latter that produced trees with average support above 50%. The ASTRAL score shows the percentage of quartet trees from input gene trees that were present in the species tree. Thus, the right way to interpret these numbers is that the 741 genes set showed less discordance.

| **number of genes** | **number of sites with maximum 25% missing data** | **Alignment size** | **includes outgroup** | **ASTRAL score** |
|---|---|---|---|---|
| 741 | 1,317,824 | 1,563,621 | yes | 87% |
| 1987 | 2,338,739 | 2,812,488 | yes | 78% |
| 2949 | 3,597,369 | 4189581 | no | |

**Table S4.** Statistics of LASTZ alignments – only assembled sequences that can be mapped to the Anna's hummingbird reference genome with a ratio of >50%.

| Sample | Identity | Aligned length |
|---|---|---|
| *Selasphorus rufus* | 0.99 | 7,435,355 |
| *Glaucis hirsuta affinis* | 0.95 | 7,238,920 |
| *Topaza pella pella* | 0.95 | 7,085,193 |
| *Florisuga mellivora* | 0.94 | 5,414,215 |
| *Archilochus colubris* | 0.99 | 7,317,584 |
| *Eutoxeres aquila* | 0.95 | 7,300,485 |
| *Threnetes leucurus* | 0.95 | 7,361,761 |
| *Eulampis jugularis* | 0.96 | 7,271,344 |
| *Heliodoxa jacula henryi* | 0.97 | 8,127,458 |
| *Phaethornis guy* | 0.95 | 8,143,601 |
| *Lampornis hemileucus* | 0.98 | 7,342,927 |
| *Panterpe insignis* | 0.98 | 7,949,639 |
| *Amazilia tzacatl* | 0.98 | 7,802,728 |
| *Chalybura urochrysia melanorrhoa* | 0.98 | 8,213,602 |
| *Eupherusa nigriventris* | 0.97 | 7,905,876 |
| *Phaetornis ruber* | 0.95 | 7,802,803 |
| *Hylocharis sapphirina* | 0.98 | 7,871,143 |
| *Polytmus theresiae* | 0.96 | 7,872,495 |
| *Campylopterus hyperythrus* | 0.98 | 8,154,833 |
| *Phaethornis bourcieri* | 0.95 | 7,739,961 |
| *Lophornis pavonina* | 0.96 | 8,071,619 |
| *Calliphlox amethystina* | 0.99 | 8,316,843 |
| *Anthracothorax nigricollis nigricollis* | 0.96 | 7,837,880 |
| *Doryfera johannae* | 0.96 | 8,231,303 |
| *Thalurania furcata* | 0.98 | 7,958,790 |
| *Trochilus polytmus* | 0.98 | 8,073,870 |
| *Heliomaster furcifer* | 0.98 | 8,004,592 |
| *Patagona gigas* | 0.98 | 7,872,679 |
| *Oreotrochilus estella* | 0.97 | 7,585,159 |
| *Microstilbon burmeisteri* | 0.99 | 8,254,909 |
| *Sappho sparganura* | 0.97 | 8,299,691 |
| *Colibri coruscans* | 0.96 | 7,625,215 |
| *Heiothryx barroti* | 0.96 | 8,212,584 |
| *Androdon aequatorialis* | 0.96 | 7,820,938 |
| *Metallura tyrianthina* | 0.97 | 8,080,857 |
| *Chalcostigma herrani* | 0.97 | 8,012,005 |
| *Boissonneaua matthewsii* | 0.97 | 8,280,820 |
| *Aglaeactis cupripennis* | 0.97 | 7,427,217 |
| *Ensifera ensifera* | 0.97 | 8,373,605 |
| *Coeligena coeligena* | 0.97 | 7,827,225 |
| *Ocreatus underwoodii* | 0.97 | 8,213,509 |
| *Heliangelus amethysticollis* | 0.97 | 7,566,599 |
| *Aglaiocercus kingi* | 0.97 | 8,169,128 |
| *Eriocnemis alinae* | 0.97 | 6,518,034 |
| *Phlogophilus hemileucurus* | 0.96 | 6,147,255 |
| *Ramphomicron microrhynchum* | 0.97 | 7,888,400 |

**Table S5.** Statistics of the LASTZ alignments against the Anna's hummingbird reference genome.

| Sample | Identity | Aligned length |
|---|---|---|
| *Selasphorus rufus* | 0.99 | 8,082,241 |
| *Glaucis hirsuta affinis* | 0.95 | 7,851,938 |
| *Topaza pella pella* | 0.95 | 7,527,640 |
| *Florisuga mellivora* | 0.94 | 7,576,079 |
| *Archilochus colubris* | 0.99 | 7,791,298 |
| *Eutoxeres aquila* | 0.95 | 8,103,870 |
| *Threnetes leucurus* | 0.95 | 7,943,349 |
| *Eulampis jugularis* | 0.95 | 7,808,689 |
| *Heliodoxa jacula henryi* | 0.97 | 8,654,411 |
| *Phaethornis guy* | 0.95 | 8,477,668 |
| *Lampornis hemileucus* | 0.98 | 8,712,950 |
| *Panterpe insignis* | 0.98 | 8,655,648 |
| *Amazilia tzacatl* | 0.98 | 8,625,527 |
| *Chalybura urochrysia melanorrhoa* | 0.98 | 8,517,027 |
| *Eupherusa nigriventris* | 0.97 | 8,680,757 |
| *Phaetornis ruber* | 0.95 | 8,515,270 |
| *Hylocharis sapphirina* | 0.98 | 8,543,232 |
| *Polytmus theresiae* | 0.96 | 8,608,241 |
| *Campylopterus hyperythrus* | 0.98 | 8,661,611 |
| *Phaethornis bourcieri* | 0.95 | 8,559,819 |
| *Lophornis pavonina* | 0.96 | 8,504,499 |
| *Calliphlox amethystina* | 0.99 | 8,615,380 |
| *Anthracothorax nigricollis nigricollis* | 0.96 | 8,523,644 |
| *Doryfera johannae* | 0.96 | 8,599,143 |
| *Thalurania furcata* | 0.98 | 8,640,418 |
| *Trochilus polytmus* | 0.98 | 8,441,286 |
| *Heliomaster furcifer* | 0.98 | 8,631,929 |
| *Patagona gigas* | 0.98 | 8,747,702 |
| *Oreotrochilus estella* | 0.97 | 8,682,249 |
| *Microstilbon burmeisteri* | 0.99 | 8,684,161 |
| *Sappho sparganura* | 0.97 | 8,501,671 |
| *Colibri coruscans* | 0.96 | 8,414,353 |
| *Heiothryx barroti* | 0.96 | 8,673,886 |
| *Androdon aequatorialis* | 0.96 | 8,582,169 |
| *Metallura tyrianthina* | 0.97 | 8,630,444 |
| *Chalcostigma herrani* | 0.97 | 8,691,487 |
| *Boissonneaua matthewsii* | 0.97 | 8,565,833 |
| *Aglaeactis cupripennis* | 0.97 | 8,732,536 |
| *Ensifera ensifera* | 0.97 | 8,580,810 |
| *Coeligena coeligena* | 0.97 | 8,653,554 |
| *Ocreatus underwoodii* | 0.97 | 8,597,582 |
| *Heliangelus amethysticollis* | 0.97 | 8,716,123 |
| *Aglaiocercus kingi* | 0.97 | 8,607,802 |
| *Eriocnemis alinae* | 0.97 | 6,661,725 |
| *Phlogophilus hemileucurus* | 0.96 | 6,274,514 |
| *Ramphomicron microrhynchum* | 0.97 | 8,687,987 |

**Table S6.** De novo assembly statistics for the 46 hummingbirds (SOAPdenovo, K=63)

| Sample | N50 | N10 | Longest | Total size |
|---|---|---|---|---|
| *Selasphorus rufus* | 251 | 455 | 2462 | 39,447,699 |
| *Glaucis hirsuta affinis* | 251 | 380 | 2606 | 68,422,247 |
| *Topaza pella pella* | 251 | 387 | 2590 | 58,544,759 |
| *Florisuga mellivora* | 251 | 367 | 1172 | 150,191,670 |
| *Archilochus colubris* | 251 | 412 | 2499 | 42,328,978 |
| *Eutoxeres aquila* | 251 | 367 | 2053 | 87,406,292 |
| *Threnetes leucurus* | 251 | 388 | 2670 | 66,773,811 |
| *Eulampis jugularis* | 251 | 413 | 2670 | 55,174,936 |
| *Heliodoxa jacula henryi* | 262 | 684 | 5449 | 32,176,045 |
| *Phaethornis guy* | 255 | 614 | 5449 | 34,519,567 |
| *Lampornis hemileucus* | 242 | 468 | 4926 | 104,000,000 |
| *Panterpe insignis* | 242 | 573 | 3673 | 48,728,973 |
| *Amazilia tzacatl* | 239 | 528 | 5193 | 61,167,084 |
| *Chalybura urochrysia melanorrhoa* | 295 | 701 | 5449 | 24,435,295 |
| *Eupherusa nigriventris* | 243 | 535 | 4354 | 65,585,920 |
| *Phaetornis ruber* | 242 | 483 | 6686 | 75,059,848 |
| *Hylocharis sapphirina* | 240 | 570 | 5613 | 33,358,586 |
| *Polytmus theresiae* | 240 | 525 | 5449 | 69,947,531 |
| *Campylopterus hyperythrus* | 248 | 645 | 6429 | 38,046,077 |
| *Phaethornis bourcieri* | 241 | 505 | 5456 | 72,940,749 |
| *Lophornis pavonina* | 243 | 583 | 5482 | 36,454,198 |
| *Calliphlox amethystina* | 291 | 700 | 5449 | 23,571,160 |
| *Anthracothorax nigricollis nigricollis* | 238 | 504 | 5703 | 60,532,736 |
| *Doryfera johannae* | 245 | 618 | 7593 | 42,696,474 |
| *Thalurania furcata* | 254 | 639 | 6314 | 27,483,735 |
| *Trochilus polytmus* | 248 | 553 | 5385 | 26,629,248 |
| *Heliomaster furcifer* | 252 | 632 | 5449 | 27,806,295 |
| *Patagona gigas* | 241 | 642 | 5677 | 48,567,911 |
| *Oreotrochilus estella* | 240 | 628 | 5449 | 37,206,172 |
| *Microstilbon burmeisteri* | 262 | 675 | 5449 | 29,271,710 |
| *Sappho sparganura* | 317 | 687 | 3937 | 20,463,617 |
| *Colibri coruscans* | 232 | 427 | 4313 | 55,278,620 |
| *Heiothryx barroti* | 244 | 567 | 4413 | 55,264,118 |
| *Androdon aequatorialis* | 238 | 464 | 5449 | 74,473,684 |
| *Metallura tyrianthina* | 239 | 523 | 4476 | 55,442,001 |
| *Chalcostigma herrani* | 249 | 626 | 5449 | 40,770,838 |
| *Boissonneaua matthewsii* | 251 | 610 | 5913 | 30,888,258 |
| *Aglaeactis cupripennis* | 241 | 487 | 8434 | 104,467,138 |
| *Ensifera ensifera* | 410 | 848 | 6714 | 19,130,811 |
| *Coeligena coeligena* | 256 | 720 | 8317 | 31,311,690 |
| *Ocreatus underwoodii* | 246 | 611 | 5449 | 37,012,388 |
| *Heliangelus amethysticollis* | 242 | 490 | 7699 | 97,638,182 |
| *Aglaiocercus kingi* | 245 | 606 | 6650 | 39,999,037 |
| *Eriocnemis alinae* | 236 | 397 | 5060 | 72,898,106 |
| *Phlogophilus hemileucurus* | 236 | 379 | 6523 | 67,095,600 |
| *Ramphomicron microrhynchum* | 241 | 540 | 5449 | 73,290,636 |

**Table S7.** First fossil occurrence of Daedalornithes in each continent used for the generation of an empirical calibration prior.

| Taxon | Specimen | Provenance | Age (Ma) | References |
|---|---|---|---|---|
| *Eocypselus vincenti* | MGUH 29278 | Denmark: Isle-of-Mors, Fur Formation | 55.8 – 54.6 | Dyke & Lindow 2009 |
| *Eocypselus rowei* | WDC-CGR-109 | USA: Green River Formation, Fossil Butte Member. | 52.0 | Ksepka *et. al* 2013 |
| *Collocalia buday* | QM F20907 | Australia: Queensland, Riversleigh, Camel Sputum Site | 18.5 – 17 | Boles 2001 |
| *Aegotheles zealandivetus* | NMNZ S.52917 | New Zealand: Otago, Home Hills Station, Manuherikia River, Bannockburn Formation | 19 – 16 | Worthy *et al.* 2022 |
| *Tachymarptis* sp. | – | South Africa: Cape Province, Varswater Formation at Langebaanweg | 5.33 – 3.6 | Manegold *et al.* 2013 |
| *Apus* sp. | NSMT PV 24548 | Japan: Aomori, Shiriya, Loc. 3 | 0.130 – 0.115 | Watanabe *et al.* 2018 |
| *Streptoprocne zonaris* | – | Brazil: Piauí, Toca da Janela da Barra de Antonião | 0.129 – 0.012 | Guérin *et al.* 1996 |

Institution/specimen acronyms: MGUH: Museum of Geological Survey of Denmark and Greenland, Copenhagen, Denmark; WDC: Wyoming Dinosaur Center, Thermopolis, WY, USA; QM: Queensland Museum, Brisbane, Australia; NMNZ: Museum of New Zealand Te Papa Tongarewa, Wellington, New Zealand; NSMT: National Museum of Nature and Science, Tsukuba, Japan.
